# Proteome-Wide Association Studies for Blood Lipids and Comparison with Transcriptome-Wide Association Studies

**DOI:** 10.1101/2023.08.17.553749

**Authors:** Daiwei Zhang, Boran Gao, Qidi Feng, Ani Manichaikul, Gina M. Peloso, Russell P. Tracy, Peter Durda, Kent D. Taylor, Yongmei Liu, W. Craig Johnson, Stacey Gabriel, Namrata Gupta, Joshua D. Smith, Francois Aguet, Kristin G. Ardlie, Thomas W. Blackwell, Robert E. Gerszten, Stephen S. Rich, Jerome I. Rotter, Laura J. Scott, Xiang Zhou, Seunggeun Lee

## Abstract

Blood lipid traits are treatable and heritable risk factors for heart disease, a leading cause of mortality worldwide. Although genome-wide association studies (GWAS) have discovered hundreds of variants associated with lipids in humans, most of the causal mechanisms of lipids remain unknown. To better understand the biological processes underlying lipid metabolism, we investigated the associations of plasma protein levels with total cholesterol (TC), triglycerides (TG), high-density lipoprotein cholesterol (HDL), and low-density lipoprotein cholesterol (LDL) in blood. We trained protein prediction models based on samples in the Multi-Ethnic Study of Atherosclerosis (MESA) and applied them to conduct proteome-wide association studies (PWAS) for lipids using the Global Lipids Genetics Consortium (GLGC) data. Of the 749 proteins tested, 42 were significantly associated with at least one lipid trait. Furthermore, we performed transcriptome-wide association studies (TWAS) for lipids using 9,714 gene expression prediction models trained on samples from peripheral blood mononuclear cells (PBMCs) in MESA and 49 tissues in the Genotype-Tissue Expression (GTEx) project. We found that although PWAS and TWAS can show different directions of associations in an individual gene, 40 out of 49 tissues showed a positive correlation between PWAS and TWAS signed p-values across all the genes, which suggests a high-level consistency between proteome-lipid associations and transcriptome-lipid associations.

## Introduction

Blood lipid levels, including levels of total cholesterol (TC), triglycerides (TG), high-density lipoprotein cholesterol (HDL), and low-density lipoprotein cholesterol (LDL), are heritable risk factors (Pilia et al., 2006) for coronary heart disease and stroke (Kannel et al., 1961; Willer & Mohlke, 2012), which are leading causes of death in the U. S. and other nations (Ahmad & Anderson, 2021; Roger et al., 2011). Genome-wide association studies (GWAS) have identified hundreds of loci that are significantly associated with at least one lipid trait in humans (Chen et al., 2013; de Vries et al., 2019; Graham et al., 2021; Hoffmann et al., 2018). Variant alleles associated with higher concentration of LDL are more abundant among subjects with coronary artery disease than those without (Willer et al., 2008). In addition, GWAS on lipids have facilitated the discovery of biological processes involved in lipoprotein metabolism (Burkhardt et al., 2010; Kozlitina et al., 2014; Musunuru et al., 2010).

Although GWAS have been successful in identifying loci associated with lipids, they only explain a small proportion of the heritability (Manolio et al., 2009), estimated to be 35% to 60% for TG, HDL, and LDL (Kathiresan et al., 2007). Moreover, most of these variants are located in non-coding regions with unclear functional roles (Willer et al., 2013). Because of population stratification and linkage disequilibrium, it is difficult to pinpoint the exact causal variants (Visscher et al., 2012). In addition, the large number of candidate variants severely limits the statistical power of GWAS (Brandes et al., 2020; Wang et al., 2016).

To boost the statistical power of GWAS and provide biologically meaningful interpretations, it is important to analyze downstream “omic” molecules, which include epigenetic, transcriptomic, and proteomic measurements, and then test their associations with phenotypes of interest. Recent multi-omic studies have elucidated the molecular mechanism of complex diseases (Arneson et al., 2017; Hasin et al., 2017; Leon-Mimila et al., 2019; Ramazzotti et al., 2018; Xiao et al., 2018). When downstream omic measurements are not available, which is true for many of the trait- and disease-based GWAS, the genetically expected omic values can be imputed using prediction models built upon omic and genetic data from a separate study (Gamazon et al., 2015; Gusev et al., 2016; Hu et al., 2019). An association test is then conducted on each gene between the GWAS trait and the imputed omic level. For example, based on imputed gene expression measurements, transcriptome-wide association studies (TWAS) (Cao et al., 2021; Wainberg et al., 2019; Zhu & Zhou, 2020) have been performed for various diseases and clinical characteristics, such as schizophrenia (Gusev et al., 2018), breast cancer (Bhattacharya et al., 2020), and structural neuroimaging traits (Zhao et al., 2021).

In addition to transcriptomics, proteomics provide further information for understanding complex diseases, since protein levels are downstream products of gene expression and can be more directly related to biological processes (A. P. Wingo et al., 2021). Compared to TWAS, fewer proteome-wide association studies (PWAS), imputation-based or not, have been performed. Existing PWAS have investigated the associations between proteins and colorectal cancer (Brandes et al., 2020), stroke (B.-S. Wu et al., 2022), Alzheimer’s disease (A. P. Wingo et al., 2021), depression (T. S. Wingo et al., 2021), post-traumatic stress disorder (T. S. Wingo et al., 2022), and other psychiatric disorders (J. Liu et al., 2021). Regarding blood lipids, although TWAS have identified hundreds of genes associated with them (Feng et al., 2021; Veturi et al., 2021; Yang et al., 2020), to the best of our knowledge, only one PWAS has been conducted for blood lipid traits (Schubert et al., 2022).

In this work, we investigated the association of blood protein abundance with blood lipid levels to identify proteins significantly associated with lipid variability. To conduct imputation-based PWAS, we trained genotype-based protein prediction models for protein levels measured from whole blood samples from the Multi-Ethnic Study of Atherosclerosis (MESA) (Bild et al., 2002; Burke et al., 2016). The prediction models were then applied to the GWAS data of the Global Lipids Genetics Consortium (GLGC) (Willer et al., 2013) to identify proteins that are significantly associated with at least one of TC, TG, HDL, and LDL. Moreover, to study the relationship between PWAS and TWAS for lipids, we conducted imputation-based TWAS for blood lipid traits using gene expression prediction models trained on samples from MESA peripheral blood mononuclear cells (PBMCs) and samples from 49 Genotype-Tissue Expression (GTEx) project tissues (Lonsdale et al., 2013). When comparing the TWAS and PWAS directions of association with lipid across all the genes on each of the 49 tissues, for most tissues, we found a positive correlation between the predicted PWAS and TWAS effects. However, for individual genes, we often observed opposite predicted PWAS and TWAS directions of effects.

## Methods

### Ethics statement

This work was approved by the Health Sciences and Behavioral Sciences Institutional Review Board of the University of Michigan (IRB ID: HUM00152975). All data in this work were collected previously and analyzed anonymously.

### Subjects

The Multi-Ethnic Study of Atherosclerosis (MESA), a part of the Trans-Omics for Precision Medicine program (TOPMed) (Kowalski et al., 2019; Taliun et al., 2021), investigates characteristics of subclinical cardiovascular diseases, i.e. those that are detected non-invasively before the onset of clinical signs and symptoms. The study aims to identify risk factors that can predict the progression of subclinical cardiovascular disease into clinically overt cardiovascular disease. The diverse, population-based sample includes 6,814 male and female subjects who are asymptomatic and aged between 45 and 84. The recruited participants consist of 38 percent White, 28 percent Black, 22 percent Hispanic, and 12 percent Asian (predominantly Chinese) individuals. In addition to genomic, transcriptomic, proteomic, and lipid data, the study also collected physiological, disease, demographic, lifestyle, and psychological factors (Bild et al., 2002; Burke et al., 2016).

### Preprocessing of MESA genotypes, proteomics, and transcriptomics

For the genotypes, we used the sequencing data from TOPMed (Kowalski et al., 2019; Taliun et al., 2021). We removed variants with minor allele frequency (MAF) of 0.05 or less among the TOPMed subjects, leaving 12,744,944 variants. Among the subjects who had genotypes, lipid levels, and demographic information, 1,438 of them were included in MESA. Samples with degrees of relatedness up to 2, as determined by KING (Manichaikul et al., 2010), were removed, which resulted in 1,403 subjects.

A total of 1,281 proteins were measured from 984 subjects. Protein levels were measured using a SOMAscan HTS Assay 1.3K for plasma proteins. The SOMAscan Assay is an aptamer-based multiplex protein assay. It measures protein levels by the number of protein-specific aptamers that successfully bind to their target protein, though some proteins may be targeted by multiple aptamers (Gold et al., 2010; Raffield et al., 2020; Schubert et al., 2022). In our analysis, targets that corresponded to multiple proteins were removed, which resulted in 1,212 proteins.

As part of the TOPMed MESA Multi-Omics project, the 984 participants were selected for proteomic measurement based on the following criteria. First, participant samples were restricted to those already included in the TOPMed Whole Genome Sequencing effort (Taliun et al., 2021). Second, the race and ethnicity reflected that of participants in the parent MESA cohort. Third, participants were chosen to maximize the amount of overlapping omic data. Fourth, a substantial proportion of participants had biospecimens from MESA Exams 1 and 5.

Among these participants, 935 individuals with protein levels had blood lipid measurements, genotypes, and covariate information. After inversely normalizing the protein levels, we computed the top 10 protein principal component (PC) scores and top 10 surrogate values (Lee et al., 2017) to detect outliers and adjust for unobserved factors that might adversely affect the analysis. Samples with p-values less than 0.001 for the chi-squared statistics of either the PC scores or the surrogate values were removed, leaving 918 samples (See Table S1 for sample characteristics). The inversely normalized protein levels were then adjusted for age, sex, self-reported race and ethnicity, usage of lipid-lowering medications, top 4 genetic PCs, and top 10 surrogate values. The residuals of the protein levels were used for the subsequent analyses.

RNA-seq was previously performed on MESA peripheral blood mononuclear cells (PBMCs) (Brown et al., 2019; Y. Liu et al., 2013). We used the reads per kilobase of transcript per million reads mapped (RPKM) of each gene in our analysis. After applying the same preprocessing pipeline as for the proteomics (i.e. sample matching, inverse normalization, outlier removal, and adjustment for the same set of covariates), we had 1,021 samples for 22,791 genes, which covered 1,167 out of the 1,212 genes in the proteomic data.

### Protein and gene expression prediction models based on MESA

We performed imputation-based PWAS for lipids by using SPrediXcan (Barbeira et al., 2018) to achieve higher statistical power. SPrediXcan builds an elastic net (Zou & Hastie, 2005) prediction model of the omic measurements of each gene using its cis-SNPs as predictors. These prediction models are then combined with external GWAS summary statistics to predict the associations between the omic levels and the phenotypes of interest. Intuitively, this approach can be understood as an association study between observed phenotypes and predicted omic levels. Figure 1(a) illustrates the workflow of SPrediXcan. In our analysis, we trained the elastic nets on the MESA data to predict the preprocessed protein levels from the cis-SNPs within a window extending one mega-base (MB) upstream and 1 MB downstream of the protein’s gene body (from the transcription starting site (TSS) to the transcription ending site (TES)). During model training, we restricted candidate predictive SNPs to those that are included in the GWAS.

**Figure 1:**
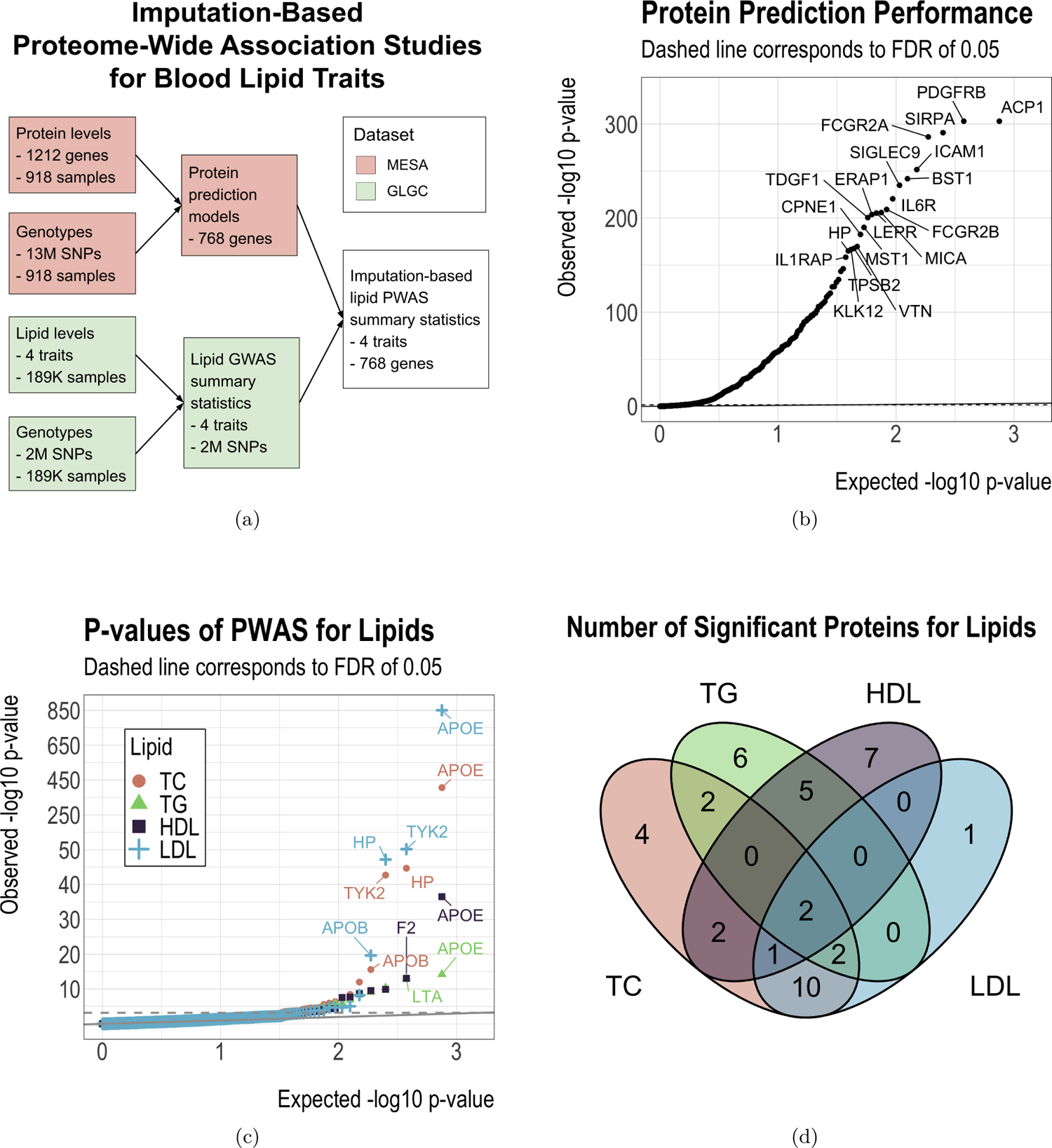
Imputation-based proteome-wide association studies (PWAS) for lipids. Panels (b) and (c): the solid line is the identity line, while the dashed line represents the false discovery rate (FDR) threshold of 0.05.

The optimal elastic net penalty weights were selected by cross-validation as recommended for SPrediXcan (Barbeira et al., 2018). We used the same procedure to build the predictive models for the transcriptomic data. After model training on the MESA data, we obtained non-trivial (i.e. at least one cis-SNP has a non-zero weight) prediction models for 749 out of 1212 proteins and 886 out of 1167 gene expressions, with an intersection of 562 genes that have both a non-trivial protein prediction model and a non-trivial gene expression prediction model.

### Gene expression prediction models based on the Genotype-Tissue Expression project

The Genotype-Tissue Expression (GTEx) project (Lonsdale et al., 2013) investigated the influence of regions in the human genome on gene expression and regulation in different tissues. Genotypes and gene expression levels were collected in 49 tissues from 900 post-mortem donors, and the sample size for each tissue ranged from 73 to 706. In our analysis, we downloaded gene expression prediction models pre-trained using the GTEx data by the authors of SPrediXcan, all of which had a predictive p-value less than 0.05. We applied the models to the GWAS summary statistics via the SPrediXcan framework to obtain tissue-specific TWAS results.

### Imputation-based PWAS and TWAS using the Global Lipids Genetics Consortium

After training the elastic nets on the MESA data, we applied the prediction models to the GWAS summary statistics from the Global Lipids Genetics Consortium (GLGC) (Willer et al., 2013). GLGC examined the associations between the genotypes and the lipid levels of 188,577 individuals of European ancestry. GWAS effect sizes and their standard errors were obtained for more than 2 million SNPs. For each blood lipid trait, we applied the protein prediction models trained on the MESA data and the tissue-specific gene expression prediction models trained on both MESA and GTEx data to the GLGC summary statistics and computed the association between the lipid and the gene’s protein and gene expression levels.

## Results

### Overview of PWAS results

Since our PWAS is imputation-based, we first assessed the prediction power of the cis-SNPs for the protein levels. Figure 1(b) shows the prediction p-values for the 749 proteins that have at least one predictive cis-SNP with a non-zero weight. The cumulative distribution function (CDF) of the predictive *r*^2^ is shown in Figure S1. With the false discovery rate (FDR) controlled at 0.05 (Ferreira & Zwinderman, 2006), 469 (63%) of the 749 proteins were significantly predictable (Figure 1 (b), Figure S1), and the predictive *r*^2^ of these proteins ranged from 0.01 to 0.80 (Figure S1). This result indicates the significance of the protein prediction models and the reliability of the imputation-based PWAS results.

We next used the protein prediction models to perform PWAS for TC, TG, HDL, LDL. The quantile-quantile plot of the PWAS p-values for each lipid is shown in Figure 1(c). Overall, we observed that 23, 17, 17, and 16 proteins were significantly associated (FDR ≤ 0.05) with TC, TG, HDL, and LDL, respectively, and 42 proteins were significantly associated with at least one lipid (Table 1, Figure 1(d)). Among these proteins, apolipoprotein E (APOE), haptoglobin (HP), and interleukin 1 receptor antagonist (IL1RN) have been identified for their associations with lipids in previous studies (Schubert et al., 2022).

**Table 1:** PWAS results for proteins that are significantly (FDR *≤* 0.05) associated with at least one blood lipid trait. Next to the PWAS (P) summary statistics of every protein, the TWAS (T) summary statistics of the same gene are also displayed. Inside each cell is the log_10_ p-value, followed by the direction of association in parentheses.

| Gene | TC |  | TG |  | HDL |  | LDL |  |
| --- | --- | --- | --- | --- | --- | --- | --- | --- |
|  | P | T | P | T | P | T | P | T |
| APOE | 406(+) | 16(-) | 14(-) | . | 37(-) | 4(+) | 850(+) | 19(-) |
| TYK2 | 43(-) | . | . | . | . | . | 53(-) | . |
| HP | 45(-) | 3(-) | 4(-) | . | . | . | 47(-) | 3(-) |
| LTA | . | . | 13(-) | . | . | . | . | . |
| MICB | 12(+) | 3(-) | 9(+) | . | . | . | 5(+) | . |
| CCL17 | . | . | . | . | 10(+) | 9(+) | . | . |
| LILRB2 | 4(-) | 4(+) | . | . | 10(-) | 8(+) | . | . |
| RBM39 | 8(-) | . | . | . | . | . | 5(-) | . |
| PCSK7 | 4(-) | 3(-) | 8(-) | 4(-) | . | . | . | . |
| FN1 | 7(+) | . | . | . | . | . | 8(+) | . |
| RSP03 | . | . | 6(+) | . | 8(-) | . | . | . |
| PDPK1 | . | . | 7(-) | . | 4(+) | . | . | . |
| MICA | 6(-) | 6(-) | . | . | 4(-) | 3(-) | 4(-) | 4(-) |
| IL1RN | 6(+) | . | . | . | . | . | 3(+) | . |
| MMP9 | . | . | 5(+) | . | 4(-) | . | . | . |
| FCGR2A | 5(-) | 6(-) | . | . | . | . | 5(-) | 6(-) |
| SERPINA1 | 4(-) | . | . | . | . | . | 5(-) | . |
| ICAM5 | 5(-) | . | . | . | . | . | 4(-) | . |
| EPHB6 | . | . | . | . | 4(-) | . | . | . |
| CTSB | . | . | 4(+) | 6(+) | . | . | . | . |
| HAVCR2 | 4(-) | . | . | . | . | . | 3(-) | . |
| MET | . | . | 4(+) | . | . | . | . | . |
| FCGR2B | 3(-) | 5(+) | . | . | . | . | 4(-) | 4(+) |
| ICAM3 | 4(+) | . | . | . | . | . | . | . |
| CPNE1 | 4(+) | 6(+) | . | . | . | . | . | . |
| COLEC11 | 4(+) | . | . | . | . | . | . | . |
| AIF1 | . | . | . | . | 4(-) | . | . | . |
| HSPA1A | . | . | 4(-) | . | . | . | . | . |
| TYRO3 | . | . | 3(+) | . | 3(-) | . | . | . |
| MMP1 | 3(-) | 3(-) | . | . | . | . | 3(-) | . |
| SHBG | . | . | . | . | 3(+) | . | . | . |
| VWF | . | . | . | . | . | . | 3(+) | . |
| AGRP | . | . | . | . | 3(+) | . | . | . |
| TKT | . | . | . | . | 3(+) | . | . | . |
| CSF3 | 4(-) | . | . | . | 8(-) | . | . | . |
| NAPA | . | . | . | . | 3(-) | . | . | . |
| APOB | 16(-) | . | 10(-) | . | 9(+) | . | 20(-) | . |
| F2 | . | . | 5(-) | . | 13(+) | . | . | . |
| HGFAC | 6(-) | . | 6(-) | . | . | . | . | . |
| MDK | 5(+) | . | . | . | . | . | . | . |
| BCAM | . | . | 3(-) | . | . | . | . | . |
| CFC1 | . | . | 4(-) | . | . | . | . | . |

### Comparison of MESA-trained PWAS and MESA-trained TWAS

To compare lipid PWAS with lipid TWAS from the same study samples, we also conducted TWAS using GLGC summary data with the predictive models trained on the MESA PBMC gene expression data. For each lipid trait, we compared the signed log p-value of the genes in PWAS and TWAS and computed the Spearman correlation coefficient (Myers & Sirois, 2006) (Figure 2), where the sign reflects the direction of association. The PWAS and TWAS signed log p-values were modestly positively correlated, where the correlation coefficient ranged from 0.083 to 0.144 and the correlation p-value were all below 0.05. For TC/TG/HDL/LDL, among the 23/17/17/16 genes whose proteins are associated with the lipid (Figure 1(d), Table 1), 10/2/4/5 genes have both protein and gene expression associated with the lipid. Out of these 10/2/4/5 genes, 6/2/2/3 genes’ protein-lipid association direction and gene expression-lipid association direction are concordant. In particular, APOE was significantly and positively associated with LDL in PWAS but significantly and negatively associated with LDL in TWAS; leukocyte immunoglobulin-like receptor B2 (LILRB2) and Fc gamma receptor IIb (FCGR2B) were significantly negatively associated with two lipids in PWAS and positively associated with the same lipids in TWAS.

**Figure 2:**
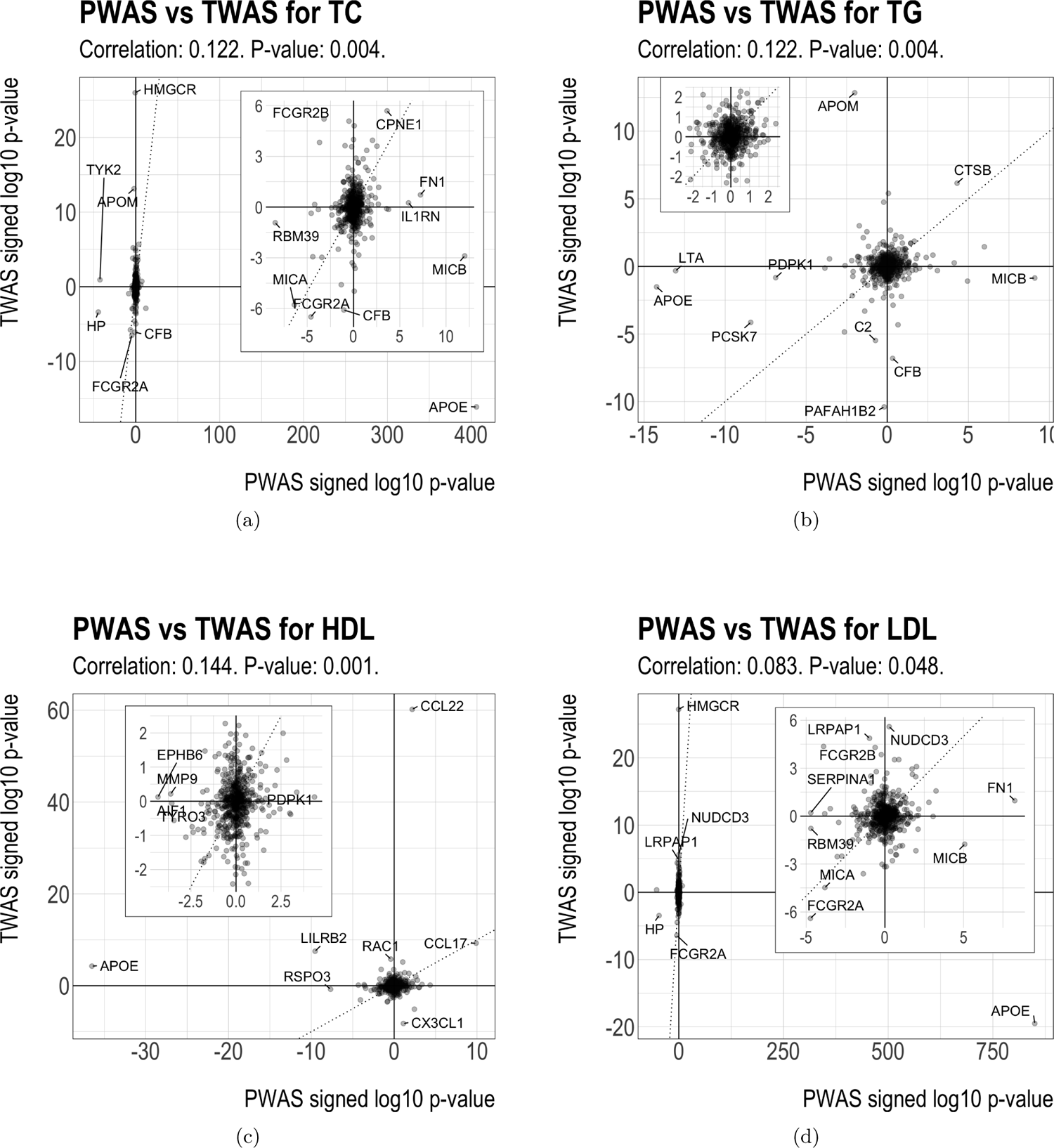
Comparison of PWAS and TWAS results for lipids. The subplot inside each panel shows zoomed results.

To better understand the opposing PWAS and TWAS effects in some of the genes, we used APOE and LDL as an example and compared the LDL GWAS summary statistics with the cis-SNPs’ weights in the protein and gene expression prediction models. Figure 3 (top panel) shows the signed log p-values of the association between LDL and the cis-SNPs of APOE in GLGC. Effect alleles were chosen so that all the GWAS effect sizes for LDL were positive.

**Figure 3:**
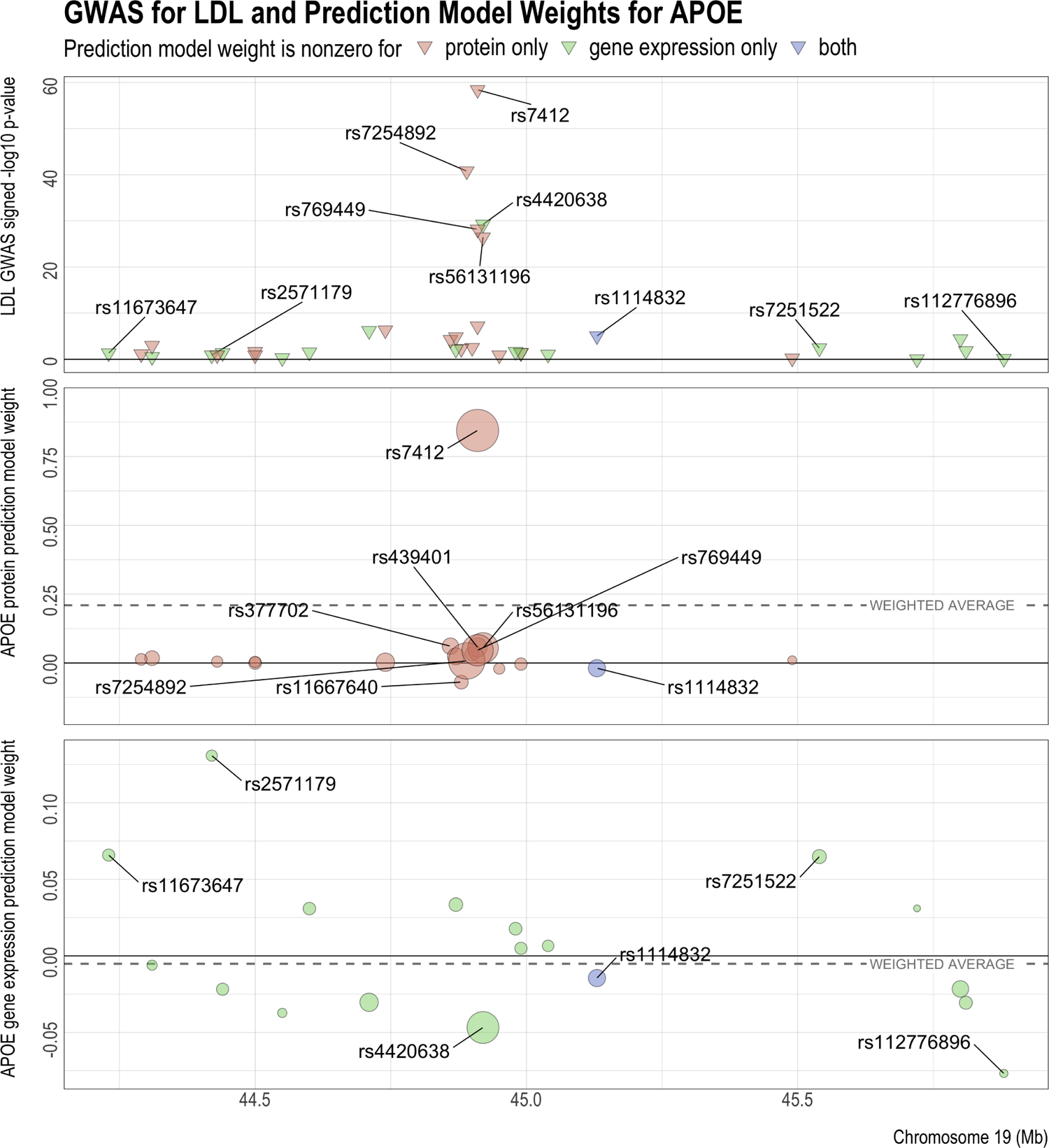
GWAS for LDL and prediction models for APOE’s protein and gene expression levels. The reference and alternative alleles for GWAS and the predictive models have been aligned and reordered so that all the SNPs have positive GWAS effects. In the center and bottom panels, the size of the circles indicates the SNP’s GWAS z-score. The z-scores are used to compute the weighted average of the model weights (dashed line), which has the same sign as and is proportional to the predicted effect of protein or gene expression on the GWAS outcome.

Among SNPs with very significant GWAS p-values, effect allele C in SNP rs7412 corresponds to the Apoε2 allele of APOE (H. Wu et al., 2020; Zhen et al., 2017). This SNP is related to the stability of the APOE isoforms (Clément-Collin et al., 2006) and is a risk factor for coronary heart disease (Tejedor et al., 2014). Another SNP with a very strong GWAS effect is rs4420638, whose effect allele G may elevate TC, TG, and HDL (Huang et al., 2015). As indicated by the colors, the sets of predictive cis-SNPs for protein and gene expression have little overlap with each other, with only one SNP (rs1114832) having a nonzero weight in both predictive models.

Figure 3 (middle panel) shows the weights of the cis-SNPs in the prediction model of APOE protein. The effects of most cis-SNPs on APOE protein had the same direction as their effects on LDL, with only four exceptions below the *y* = 0 line. In particular, the effects of rs7412 for LDL and APOE protein were both strong and of the same sign, dominating all the other cis-SNPs. Thus, the resulting association between APOE protein and LDL was positive, as indicated by the positive weighted average of the predictive weights (dashed line). On the other hand, compared to the PWAS results, the directions of the effects of the predictive cis-SNPs on APOE gene expression were approximately equally split between positive and negative, as shown in Figure 3 (bottom panel). Nevertheless, the negative weights outweighed the positive weights, with the greatest contribution from rs4420638 and rs112776896, which has a strong positive association with LDL, but strong negative association with APOE gene expression. Thus the resulting association between LDL and APOE gene expression was negative, as indicated by the negative weighted average of the gene expression predictive weights (dashed line). Overall, due to the small proportion of overlapping nonzero predictive weights and their different directions of effects (Figure S2), APOE protein and gene expression have opposite directions of association with LDL. In addition, similar patterns were observed for LDL with other genes, such as FCGR2B, LILRB2, major histocompatibility complex class I polypeptide-related sequence B (MICB) (Figures S3-S8), as well as for the other lipids (Figures S9-S16, S18-S25, S27-S34).

### Comparison of MESA-trained PWAS and GTEx-trained TWAS

The TWAS results obtained from MESA only used gene expression measurements in PBMCs. Since the gene expression levels in some tissues, such as liver, may be more relevant to lipid levels compared to those in other tissues, we extended our TWAS analysis using gene expression data from 49 GTEx tissues. Results of MESA-trained PWAS, MESA-trained TWAS, and GTEx-trained TWAS are compared in Figures 4(a), S17 (a), S26 (a), S35 (a). Overall, for all lipids, the significance and direction of association for PWAS and TWAS are heterogeneous across individual genes. For some genes, the predicted protein and gene expression levels had very consistent directions of association with LDL. For example, for major histocompatibility complex class I polypeptide-related sequence A (MICA), LDL was positively associated with both protein and gene expression in MESA and with gene expression in 43 out of 49 tissues in GTEx. Other examples with similar patterns were observed for MICA with TC and HDL, copine 1 (CPNE1) with TC, and cathepsin B (CTSB) with TG. On the other hand, for some other genes, the protein and gene expression had mixed directions of association. For instance, LDL was positively associated with HP protein levels, but had approximately equal numbers of positive and negative associations with gene expression levels across tissues. Similar inconsistent patterns were observed for HP with TC, APOE with TC and LDL, and apolipoprotein B (APOB) with TC, TG, and HDL.

**Figure 4:**
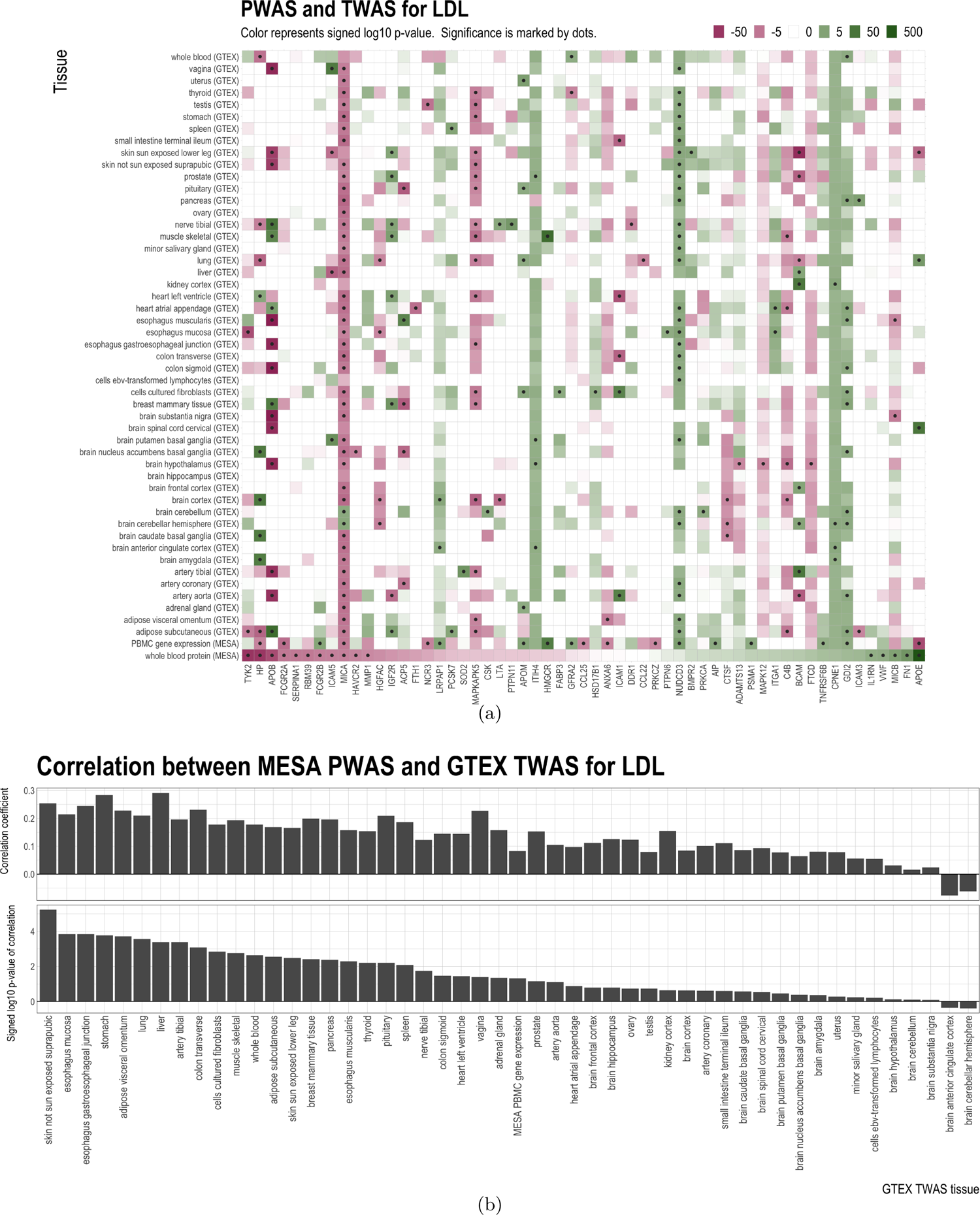
Comparison of MESA PBMC PWAS, MESA PBMC TWAS, and GTEx tissue-specific TWAS results for LDL. Panel (a): signed log p-value and significance of association. Missing values are shown in white. Significance of association is deteremined by the false discovery rate (FDR) threshold of 0.05. Only genes with at least one significant association with LDL are displayed. Panel (b): correlation between signed log p-values of MESA PBMC PWAS and signed log p-values of each GTEx tissue-specific TWAS (i.e. the correlation between the bottom row and every other row of the grid in Panel (a)).

We next evaluated the correlation patterns of PWAS and TWAS effects when aggregated across all the genes and how this correlation varied across tissues. Figure 4(b) shows the Spearman correlations for each tissue between the signed log p-values for MESA-trained PWAS and GTEx-trained TWAS for LDL. Out of the 49 tissues in GTEx, the PWAS-TWAS correlation was positive in 47 of them (binomial test p-value: 2.2 × 10^-12^). For TC, TG, and HDL, the PWAS-TWAS correlations were positive in 41, 43, 40 tissues, respectively (Figures S17(b), S26(b), and 35(b)). These findings indicate that although the relation between the proteins’ and the tissue-specific gene expressions’ effects on lipids can be mixed on a single gene, the aggregated correlations between TWAS and PWAS results for lipids across all genes were mostly positive, even if the gene expression predictive models and the protein predictive models were trained using different datasets (i.e. MESA and GTEx).

## Discussion

In this work, we conducted PWAS for blood lipids and identified 42 proteins significantly associated with at least one of TC, TG, HDL, and LDL. Several of these proteins, such as tyrosine kinase 2 (TYK2) (Grunert et al., 2011; Qi et al., 2019), MICA and MICB (Bilotta et al. 2019; Yamamoto et al. 2001), IL1RN (Schubert et al., 2022), HP (Braeckman et al., 1999; Schubert et al., 2022), APOE and APOB (Abd El-Aziz & Mohamed, 2016; Schubert et al., 2022; The Emerging Risk Factors Collaboration*, 2009; Weisgraber, 1994), have been previously identified for their association with blood lipids and related diseases. In particular, we found APOE and APOB to be significantly associated with all four lipid traits. Other proteins, such as lymphotoxin alpha (LTA), C-C motif chemokine ligand 17 (CCCL17), and LILRB2, are novel proteins that have not been previously identified for their associations with blood lipids.

Moreover, we conducted TWAS for blood lipids in different tissues and compared the results with the PWAS results. We found that PWAS and TWAS effects for lipids were heterogeneous across tissues and genes, and demonstrated that one cause of this discrepancy is the limited proportion of overlapping SNPs with nonzero predictive weights and their different directions of effect. Nevertheless, when we computed the correlation between the PWAS and TWAS signed log p-values for all the genes in every tissue, the correlation coefficients across various tissues were almost all positive. These results demonstrate that for a single gene, its gene expression’s association with lipids may differ from its protein’s association with lipids, but when the results for all the genes are aggregated, the lipid TWAS and lipid PWAS results are more consistent.

One limitation of our analyses is that not all confounders of omic or lipid levels might have been accounted for. Blood lipids in GWAS can come from a variety of sources, and there could be factors that are correlated with omic levels but not included in the study. Similarly, for training the omic prediction models, although we computed the surrogate values to adjust for unobserved factors that are relevant to the analysis, there could still be factors that are not reflected by the surrogate values and other covariates in the model, such as those related to the collection, processing, and storage of blood or plasma as well as machine artifacts. Furthermore, the set of covariates included in the GWAS might not be the same as those that are adjusted for in the omic prediction models. These potential issues with the covariates and unobserved factors may cause suboptimal accuracy or efficiency in the imputation-based PWAS and TWAS results.

A limitation of our tissue-specific GTEx-based TWAS for lipids is the high number of missing gene-tissue pairs, due to their absence in the GTEx data. Imputation methods can be applied to these gene-tissue pairs, so that the missing signed p-values of the tissue-specific gene expression-lipid associations could be imputed, which could provide more insight into the connection between the lipid PWAS and lipid TWAS.

Another limitation of our analyses is that for training the omic prediction models, samples from all ancestry groups were used in order to gain power, but in GLGC, most samples are European. This discrepancy in study populations could cause inaccuracy in the analysis (Abdellaoui et al., 2019; Price et al., 2010; Zhang et al., 2020). A multi-ethnic omic dataset with a larger sampler size than MESA will facilitate the training of ancestry-specific, high-power prediction models, and lipid GWAS with more diverse samples will make imputation–based lipid PWAS and lipid TWAS findings more applicable to individuals from non-European populations (Bhattacharya et al., 2020; Keys et al., 2020).

## Acknowledgements

This research was mainly supported by NIH grant R01HL142023. Whole genome sequencing (WGS) for the Trans-Omics in Precision Medicine (TOPMed) program was supported by the National Heart, Lung and Blood Institute (NHLBI). WGS for “NHLBI TOPMed: Multi-Ethnic Study of Atherosclerosis (MESA)” (phs001416.v1.p1) was performed at the Broad Institute of MIT and Harvard (3U54HG003067-13S1). Centralized read mapping and genotype calling, along with variant quality metrics and filtering were provided by the TOPMed Informatics Research Center (3R01HL-117626-02S1). Phenotype harmonization, data management, sample-identity QC, and general study coordination, were provided by the TOPMed Data Coordinating Center (3R01HL-120393-02S1), and TOPMed MESA Multi-Omics (HHSN2682015000031/HSN26800004). The MESA projects are conducted and supported by the National Heart, Lung, and Blood Institute (NHLBI) in collaboration with MESA investigators. Support for the Multi-Ethnic Study of Atherosclerosis (MESA) projects are conducted and supported by the National Heart, Lung, and Blood Institute (NHLBI) in collaboration with MESA investigators. Support for MESA is provided by contracts 75N92020D00001, HHSN268201500003I, N01-HC-95159, 75N92020D00005, N01-HC-95160, 75N92020D00002, N01-HC-95161, 75N92020D00003, N01-HC-95162, 75N92020D00006, N01-HC-95163, 75N92020D00004, N01-HC-95164, 75N92020D00007, N01-HC-95165, N01-HC-95166, N01-HC-95167, N01-HC-95168, N01-HC-95169, UL1-TR-000040, UL1-TR-001079, UL1-TR-001420, UL1TR001881, DK063491, and R01HL105756. The authors thank the other investigators, the staff, and the participants of the MESA study for their valuable contributions. A full list of participating MESA investigators and institutes can be found at http://www.mesa-nhlbi.org. SL is supported by by the Brain Pool Plus (BP+, Brain Pool+) Program through the National Research Foundation of Korea (NRF) funded by the Ministry of Science and ICT (2020H1D3A2A03100666). GMP is supported by NIH grants R01HL142711 and R01HL127564. The authors also thank Hae Kyung Im and Alvaro Barbeira for their help with using SPrediXcan.

## Notes

https://upenn.box.com/s/8hvds35u5gmfbxuyll0nwxznyi1wj288

### Competing Interest Statement

The authors have declared no competing interest.

