## Supplementary Information for "Proteome-Wide Association Studies for Blood Lipids and Comparison with Transcriptome-Wide Association Studies"

### Supplementary Materials

### S1 Additional results for all lipids

Table S1: Data characteristics of the MESA dataset. For continuous variables, the mean and standard deviation (in parentheses) are displayed.

| Self-reported race | Asian (7%) | Black (20%) | Hispanic (31%) | White (43%) |
|---:|---:|---:|---:|---:|
| TC (mg/dl) | 196 (29) | 189 (39) | 197 (35) | 197 (33) |
| TG (mg/dl) | 150 (74) | 93 (42) | 145 (66) | 126 (63) |
| HDL (mg/dl) | 49 (11) | 52 (14) | 48 (12) | 53 (15) |
| LDL (mg/dl) | 117 (26) | 118 (34) | 119 (33) | 119 (30) |
| Age (mg/dl) | 62 (10) | 61 (10) | 59 (09) | 61 (10) |
| Sex (female) | 45% | 59 % | 53 % | 52 % |
| Using lipid medication | 22% | 13 % | 12 % | 19 % |

Figure S1: MESA protein prediction model performance.

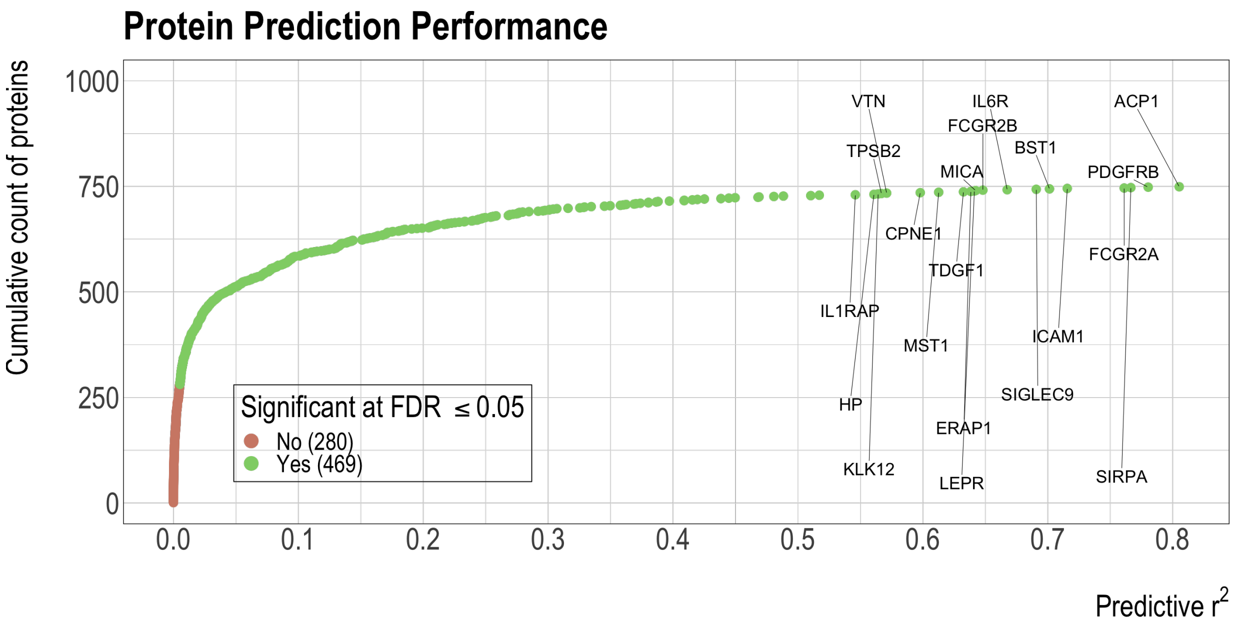

#### S2 Additional results for low-density lipoprotein (LDL)

Figure S2: Comparison of APOE's protein and gene expression predictive model weights with the LDL GWAS z-scores of the SNPs. The reference and alternative alleles for GWAS and the predictive models have been aligned and reordered so that all the SNPs have positive GWAS effects. The z-scores are used to compute the weighted average of the model weights (dashed lines), which have the same signs as and are proportional to the predicted effects of protein and gene expression on the GWAS outcome.

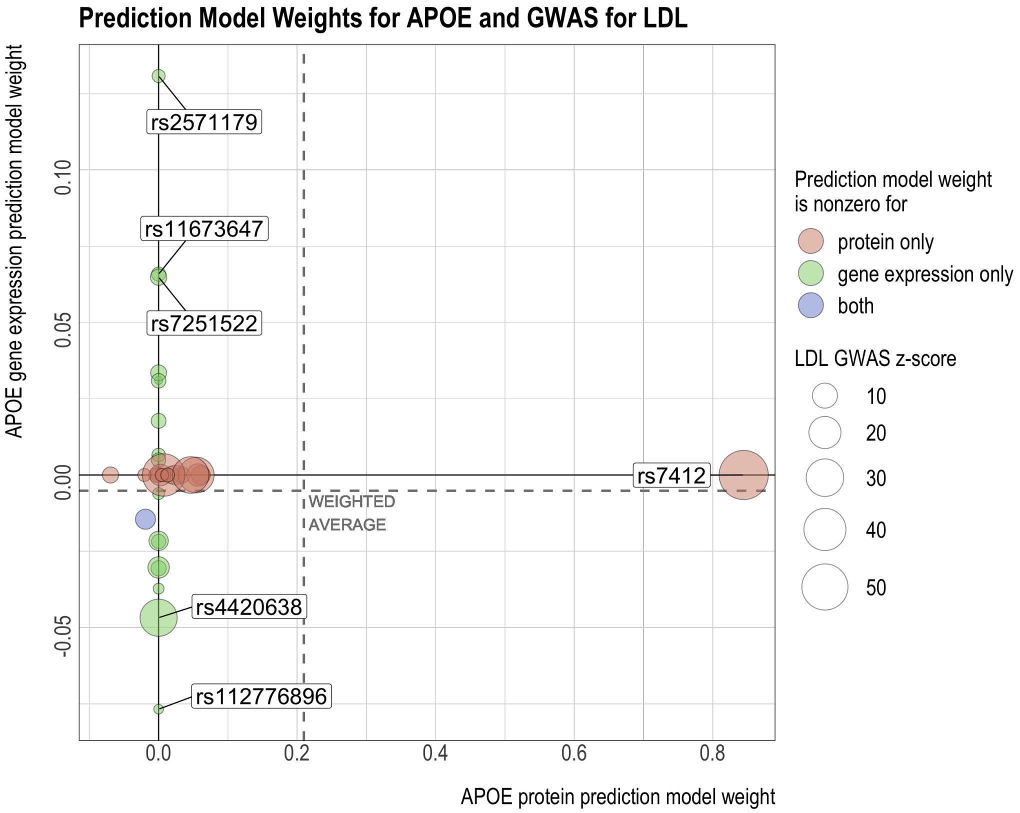

Figure S3: GWAS for LDL and prediction models for FCGR2B's protein and gene expression levels. The reference and alternative alleles for GWAS and the predictive models have been aligned and reordered so that all the SNPs have positive GWAS effects. In the center and bottom panels, the size of the circles indicates the SNP's GWAS z-score. The z-scores are used to compute the weighted average of the model weights (dashed line), which has the same sign as and is proportional to the predicted effect of protein or gene expression on the GWAS outcome.

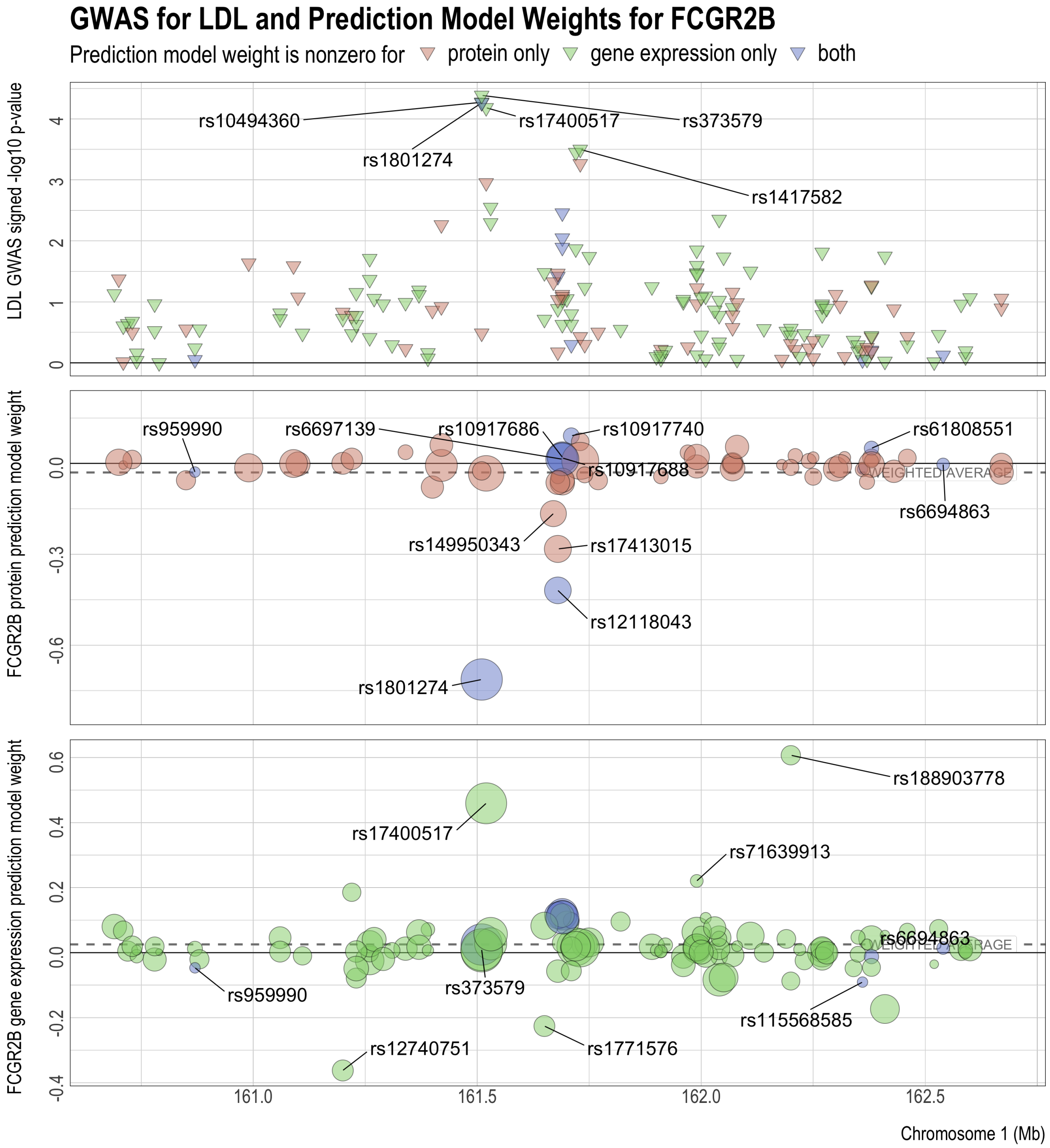

Figure S4: Comparison of FCGR2B's protein and gene expression predictive model weights with the LDL GWAS z-scores of the SNPs. The reference and alternative alleles for GWAS and the predictive models have been aligned and reordered so that all the SNPs have positive GWAS effects. The z-scores are used to compute the weighted average of the model weights (dashed lines), which have the same signs as and are proportional to the predicted effects of protein and gene expression on the GWAS outcome.

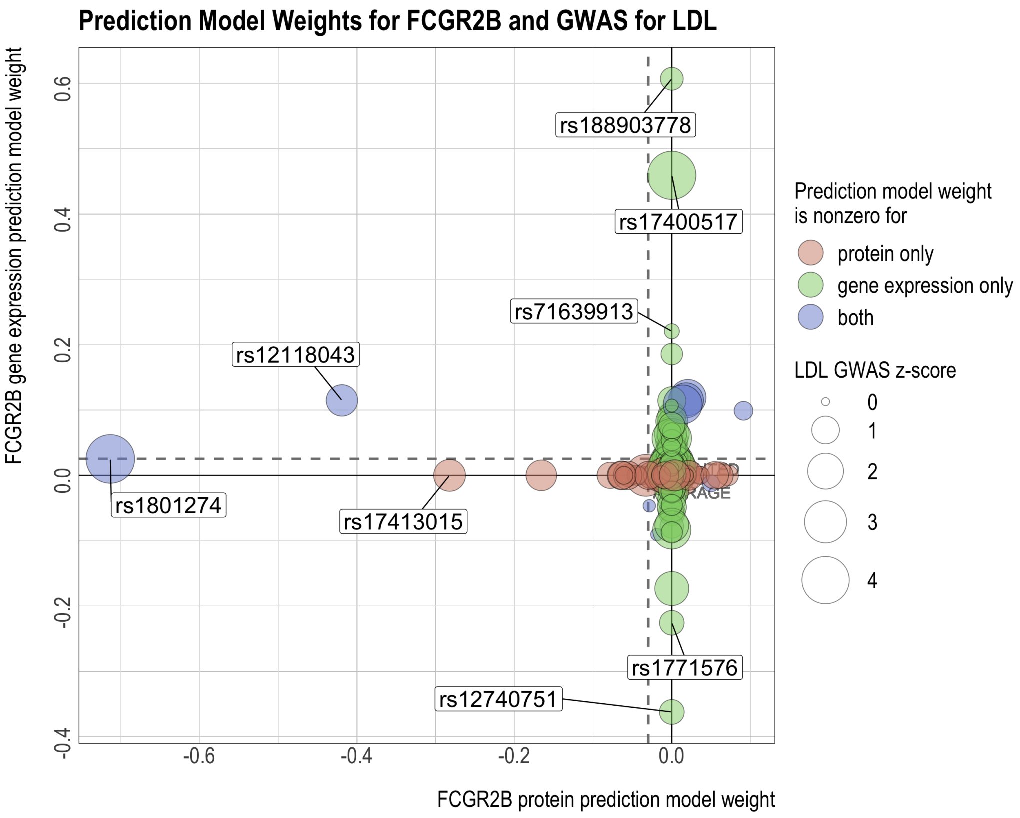

Figure S5: GWAS for LDL and prediction models for LILRB2's protein and gene expression levels. The reference and alternative alleles for GWAS and the predictive models have been aligned and reordered so that all the SNPs have positive GWAS effects. In the center and bottom panels, the size of the circles indicates the SNP's GWAS z-score. The z-scores are used to compute the weighted average of the model weights (dashed line), which has the same sign as and is proportional to the predicted effect of protein or gene expression on the GWAS outcome.

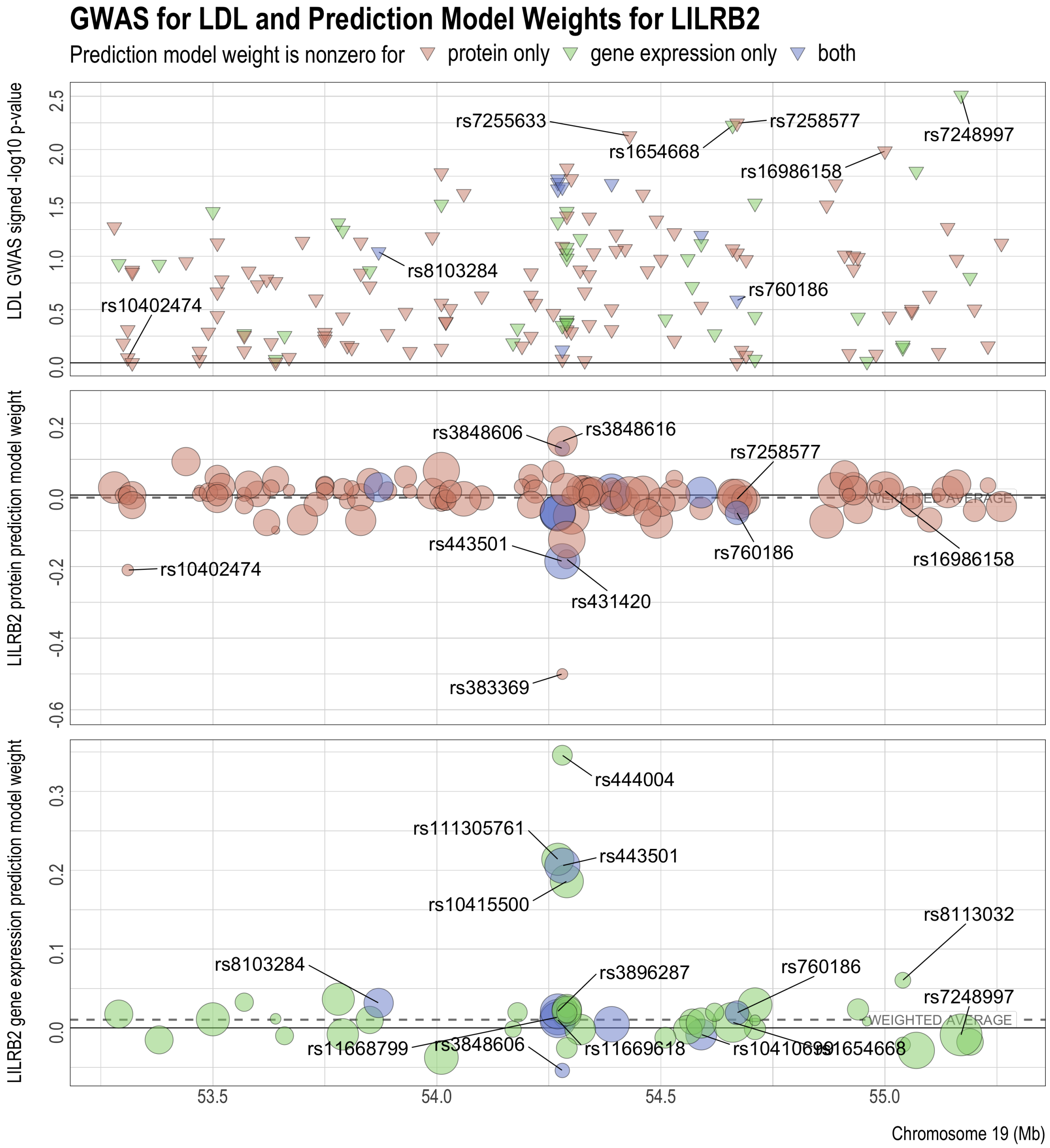

Figure S6: Comparison of LILRB2's protein and gene expression predictive model weights with the LDL GWAS z-scores of the SNPs. The reference and alternative alleles for GWAS and the predictive models have been aligned and reordered so that all the SNPs have positive GWAS effects. The z-scores are used to compute the weighted average of the model weights (dashed lines), which have the same signs as and are proportional to the predicted effects of protein and gene expression on the GWAS outcome.

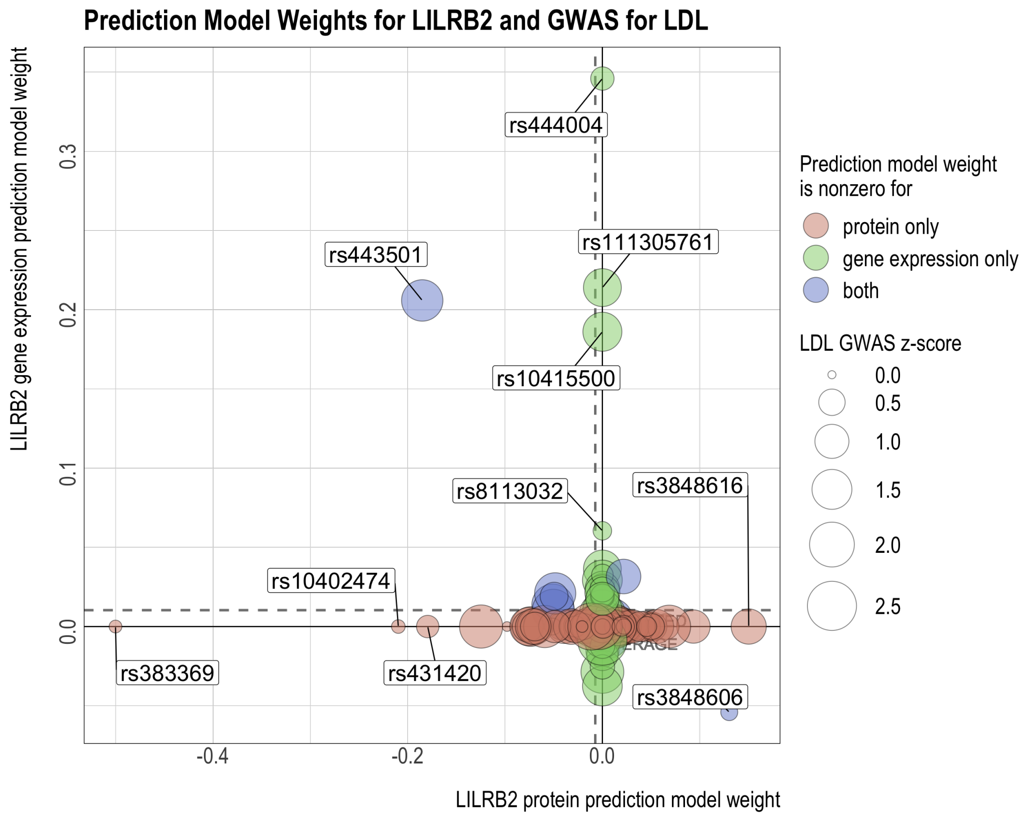

Figure S7: GWAS for LDL and prediction models for MICB's protein and gene expression levels. The reference and alternative alleles for GWAS and the predictive models have been aligned and reordered so that all the SNPs have positive GWAS effects. In the center and bottom panels, the size of the circles indicates the SNP's GWAS z-score. The z-scores are used to compute the weighted average of the model weights (dashed line), which has the same sign as and is proportional to the predicted effect of protein or gene expression on the GWAS outcome.

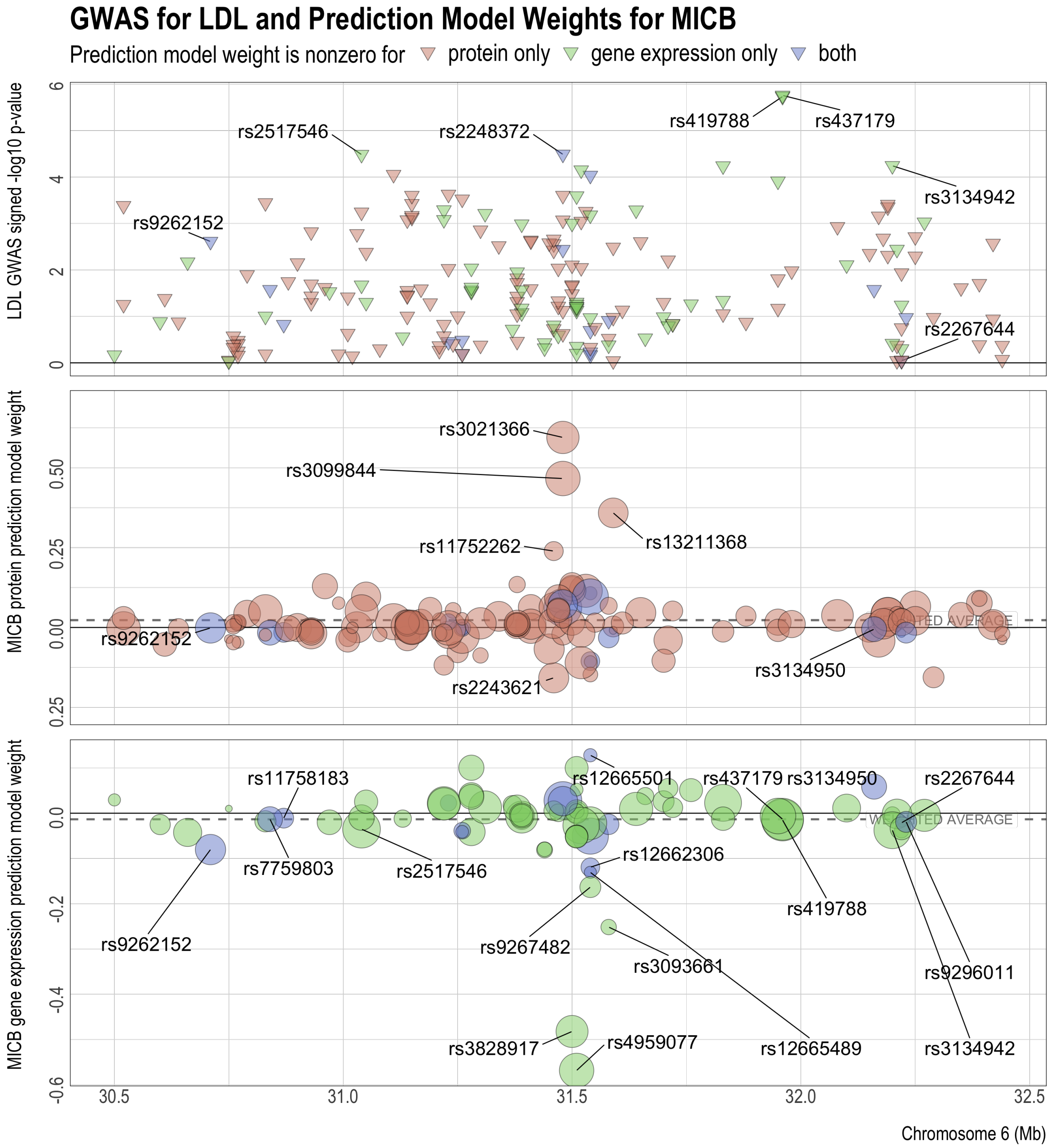

Figure S8: Comparison of MICB's protein and gene expression predictive model weights with the LDL GWAS z-scores of the SNPs. The reference and alternative alleles for GWAS and the predictive models have been aligned and reordered so that all the SNPs have positive GWAS effects. The z-scores are used to compute the weighted average of the model weights (dashed lines), which have the same signs as and are proportional to the predicted effects of protein and gene expression on the GWAS outcome.

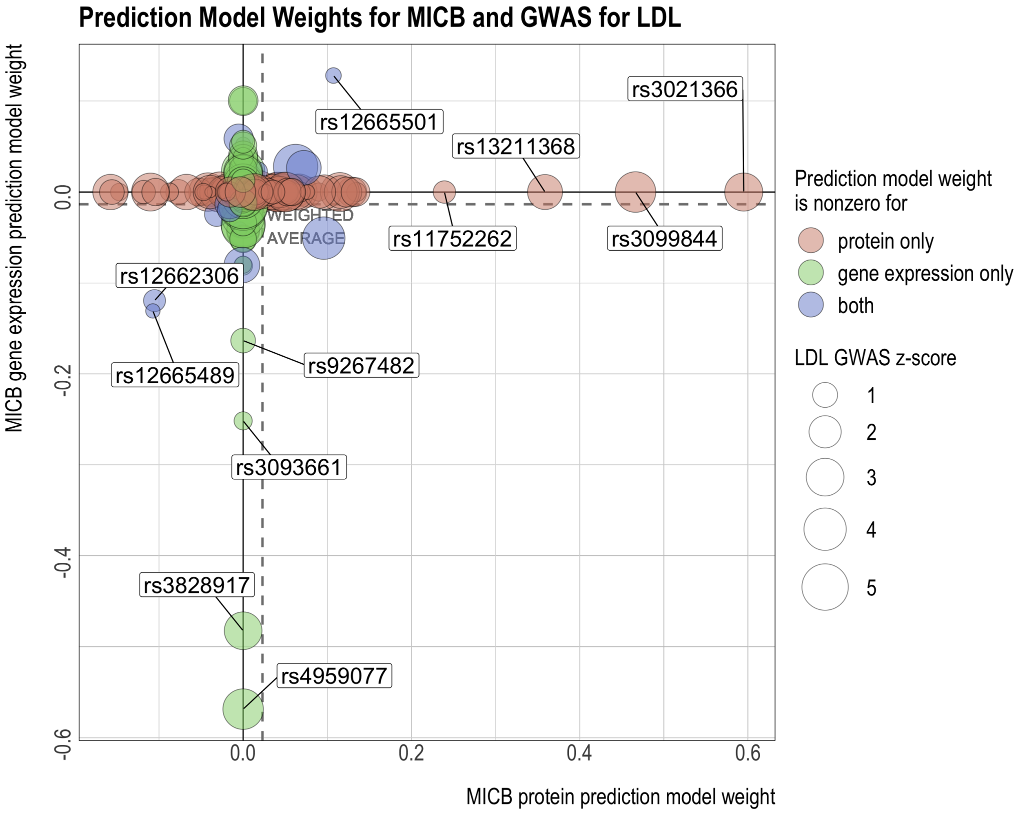

### S3 Additional results for total cholesterol (TC)

Figure S9: GWAS for TC and prediction models for APOE's protein and gene expression levels. The reference and alternative alleles for GWAS and the predictive models have been aligned and reordered so that all the SNPs have positive GWAS effects. In the center and bottom panels, the size of the circles indicates the SNP's GWAS z-score. The z-scores are used to compute the weighted average of the model weights (dashed line), which has the same sign as and is proportional to the predicted effect of protein or gene expression on the GWAS outcome.

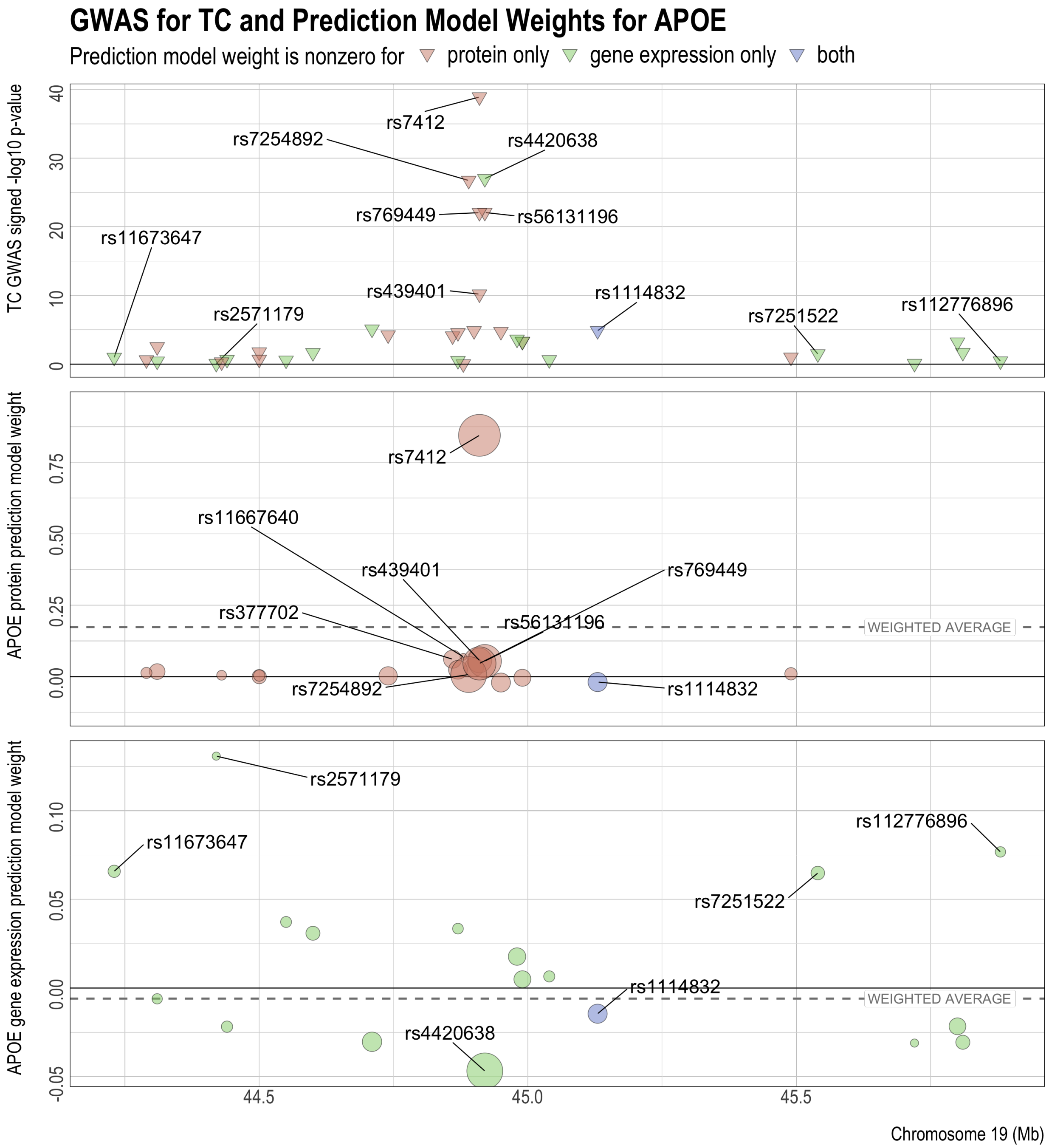

Figure S10: Comparison of APOE's protein and gene expression predictive model weights with the TC GWAS z-scores of the SNPs. The reference and alternative alleles for GWAS and the predictive models have been aligned and reordered so that all the SNPs have positive GWAS effects. The z-scores are used to compute the weighted average of the model weights (dashed lines), which have the same signs as and are proportional to the predicted effects of protein and gene expression on the GWAS outcome.

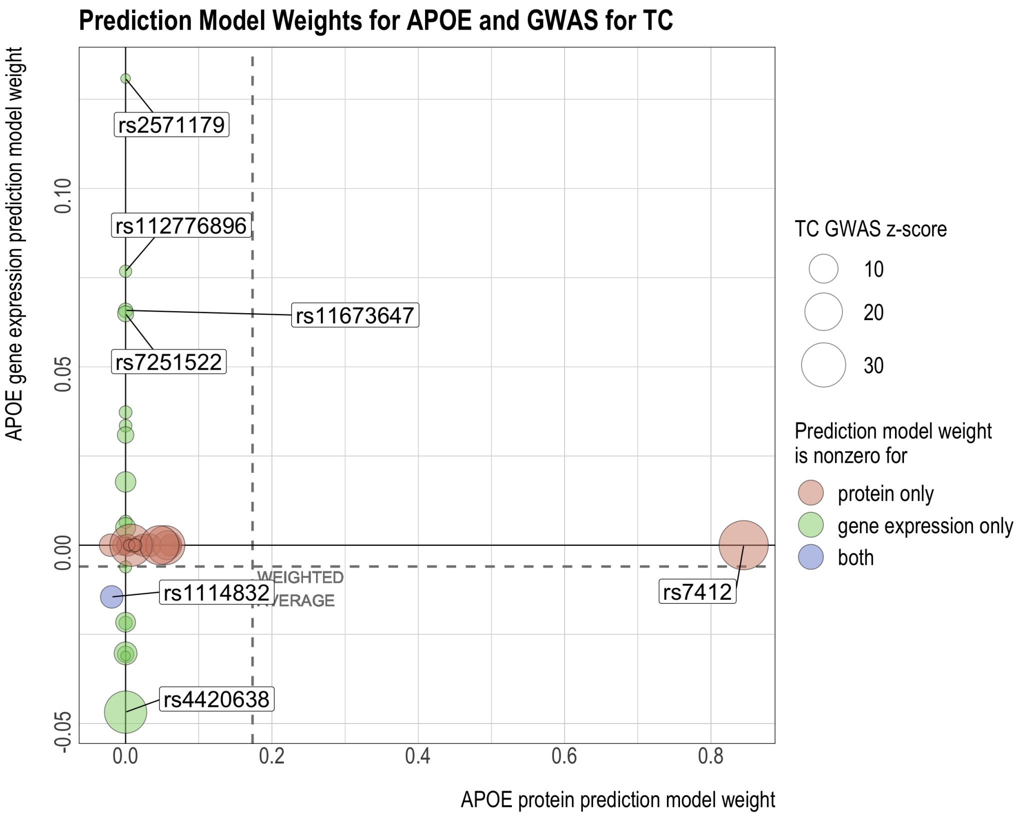

Figure S11: GWAS for TC and prediction models for FCGR2B's protein and gene expression levels. The reference and alternative alleles for GWAS and the predictive models have been aligned and reordered so that all the SNPs have positive GWAS effects. In the center and bottom panels, the size of the circles indicates the SNP's GWAS z-score. The z-scores are used to compute the weighted average of the model weights (dashed line), which has the same sign as and is proportional to the predicted effect of protein or gene expression on the GWAS outcome.

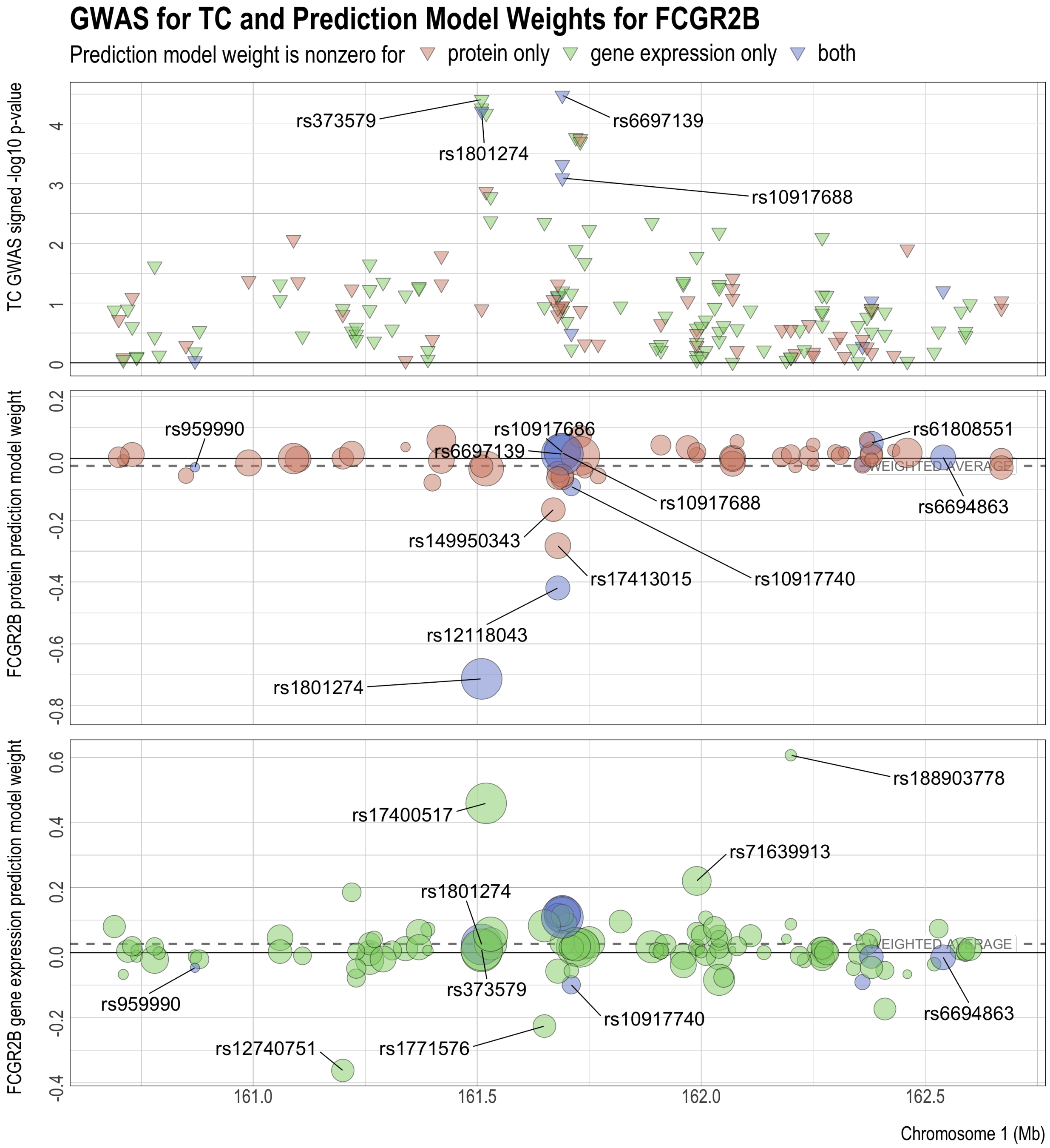

Figure S12: Comparison of FCGR2B's protein and gene expression predictive model weights with the TC GWAS z-scores of the SNPs. The reference and alternative alleles for GWAS and the predictive models have been aligned and reordered so that all the SNPs have positive GWAS effects. The z-scores are used to compute the weighted average of the model weights (dashed lines), which have the same signs as and are proportional to the predicted effects of protein and gene expression on the GWAS outcome.

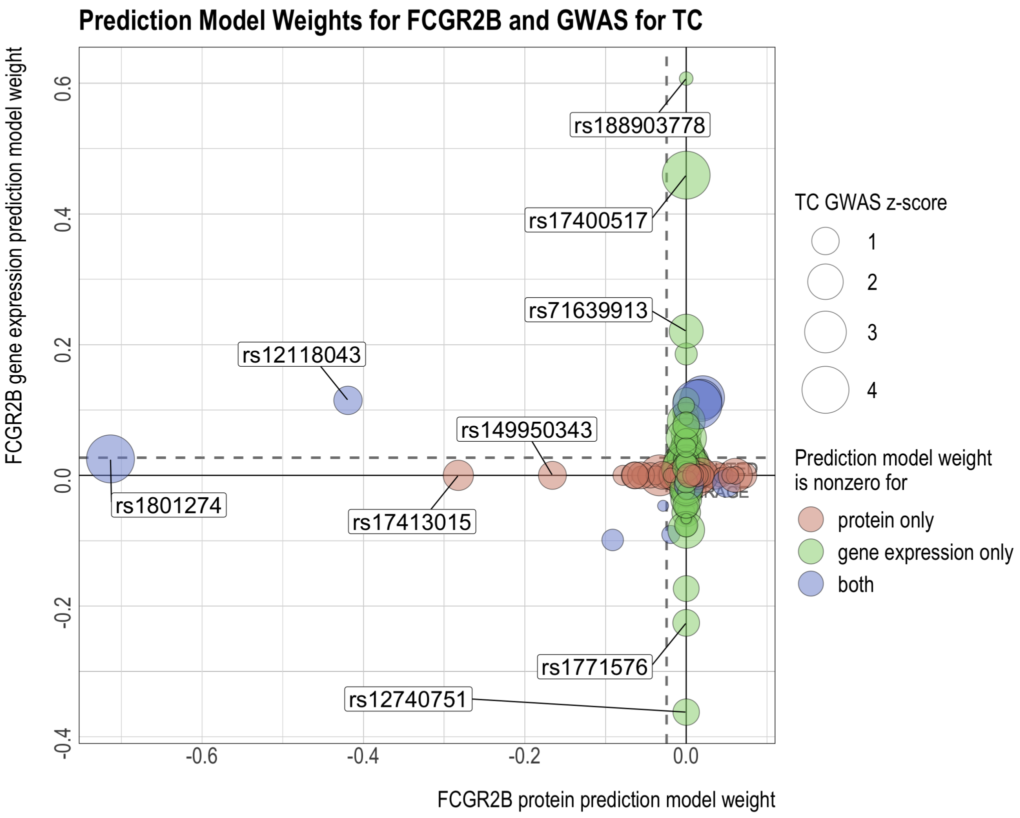

Figure S13: GWAS for TC and prediction models for LILRB2's protein and gene expression levels. The reference and alternative alleles for GWAS and the predictive models have been aligned and reordered so that all the SNPs have positive GWAS effects. In the center and bottom panels, the size of the circles indicates the SNP's GWAS z-score. The z-scores are used to compute the weighted average of the model weights (dashed line), which has the same sign as and is proportional to the predicted effect of protein or gene expression on the GWAS outcome.

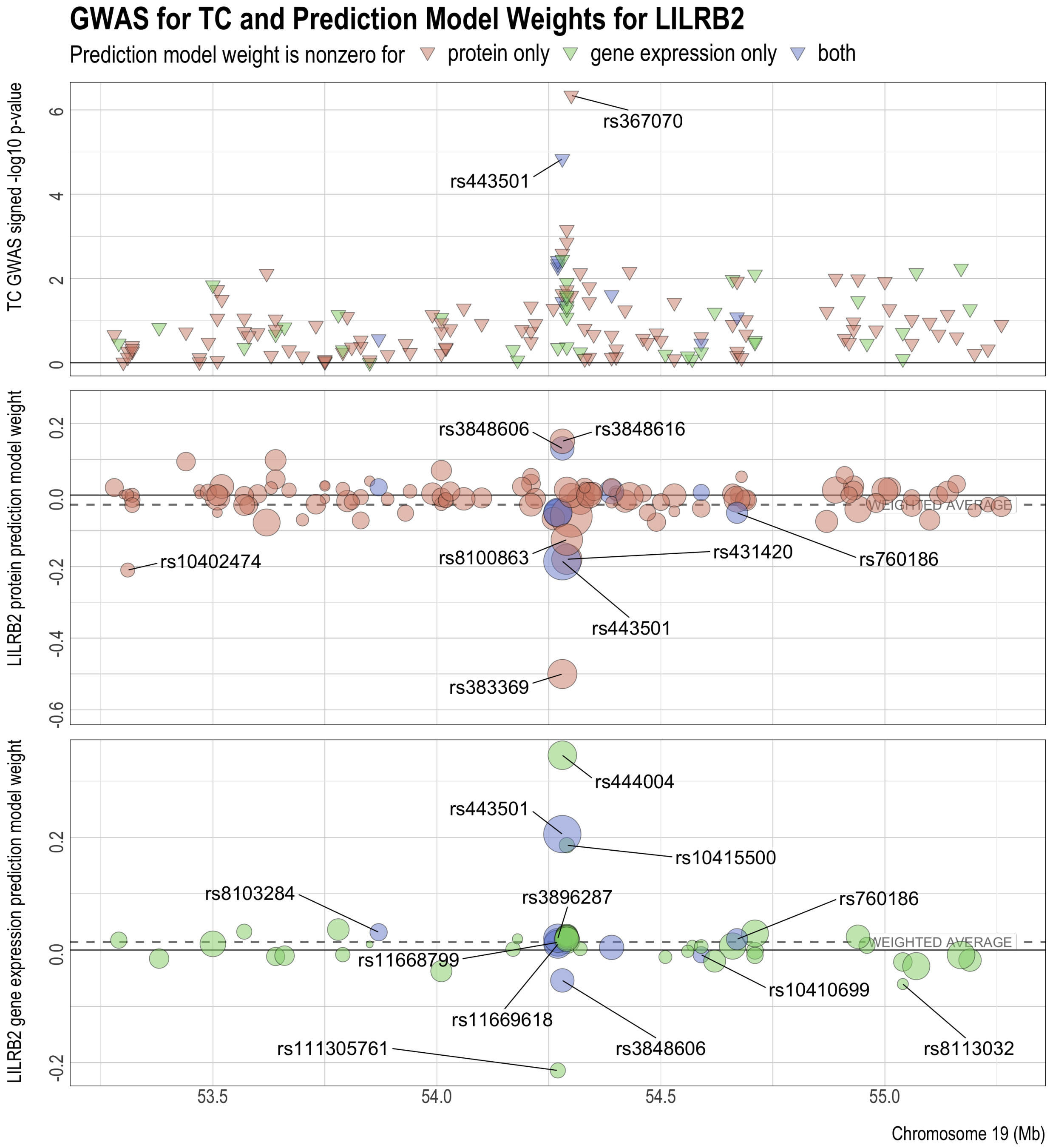

Figure S14: Comparison of LILRB2's protein and gene expression predictive model weights with the TC GWAS z-scores of the SNPs. The reference and alternative alleles for GWAS and the predictive models have been aligned and reordered so that all the SNPs have positive GWAS effects. The z-scores are used to compute the weighted average of the model weights (dashed lines), which have the same signs as and are proportional to the predicted effects of protein and gene expression on the GWAS outcome.

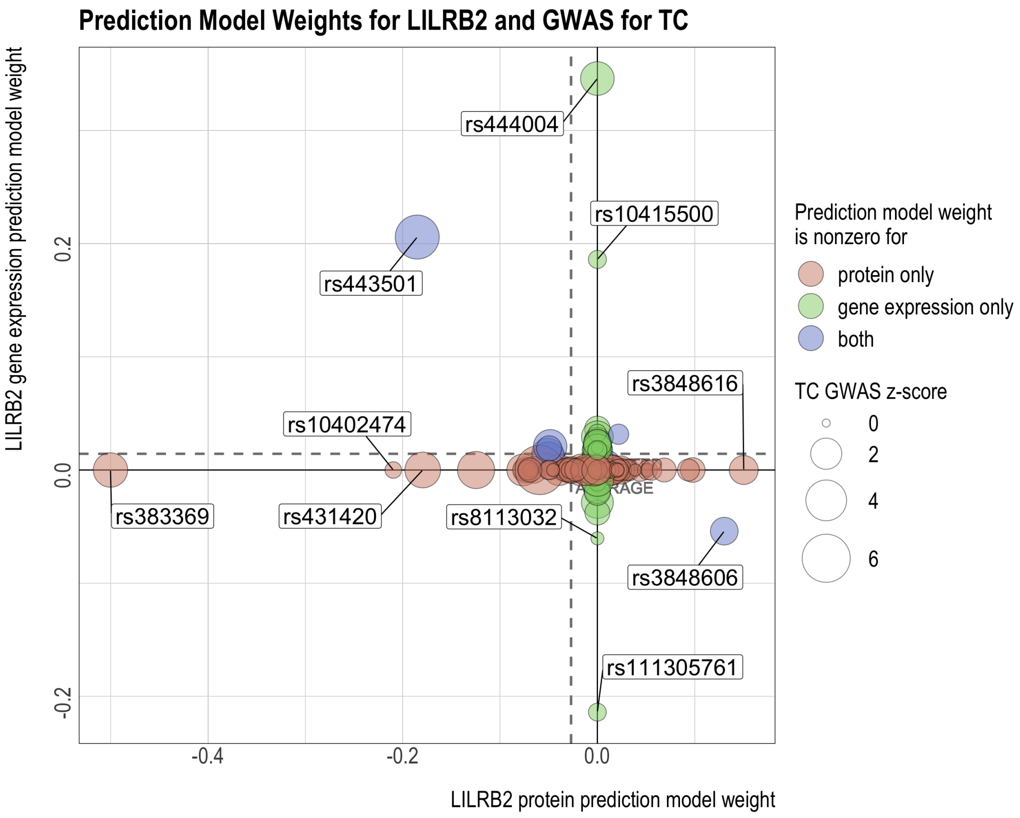

Figure S15: GWAS for TC and prediction models for MICB's protein and gene expression levels. The reference and alternative alleles for GWAS and the predictive models have been aligned and reordered so that all the SNPs have positive GWAS effects. In the center and bottom panels, the size of the circles indicates the SNP's GWAS z-score. The z-scores are used to compute the weighted average of the model weights (dashed line), which has the same sign as and is proportional to the predicted effect of protein or gene expression on the GWAS outcome.

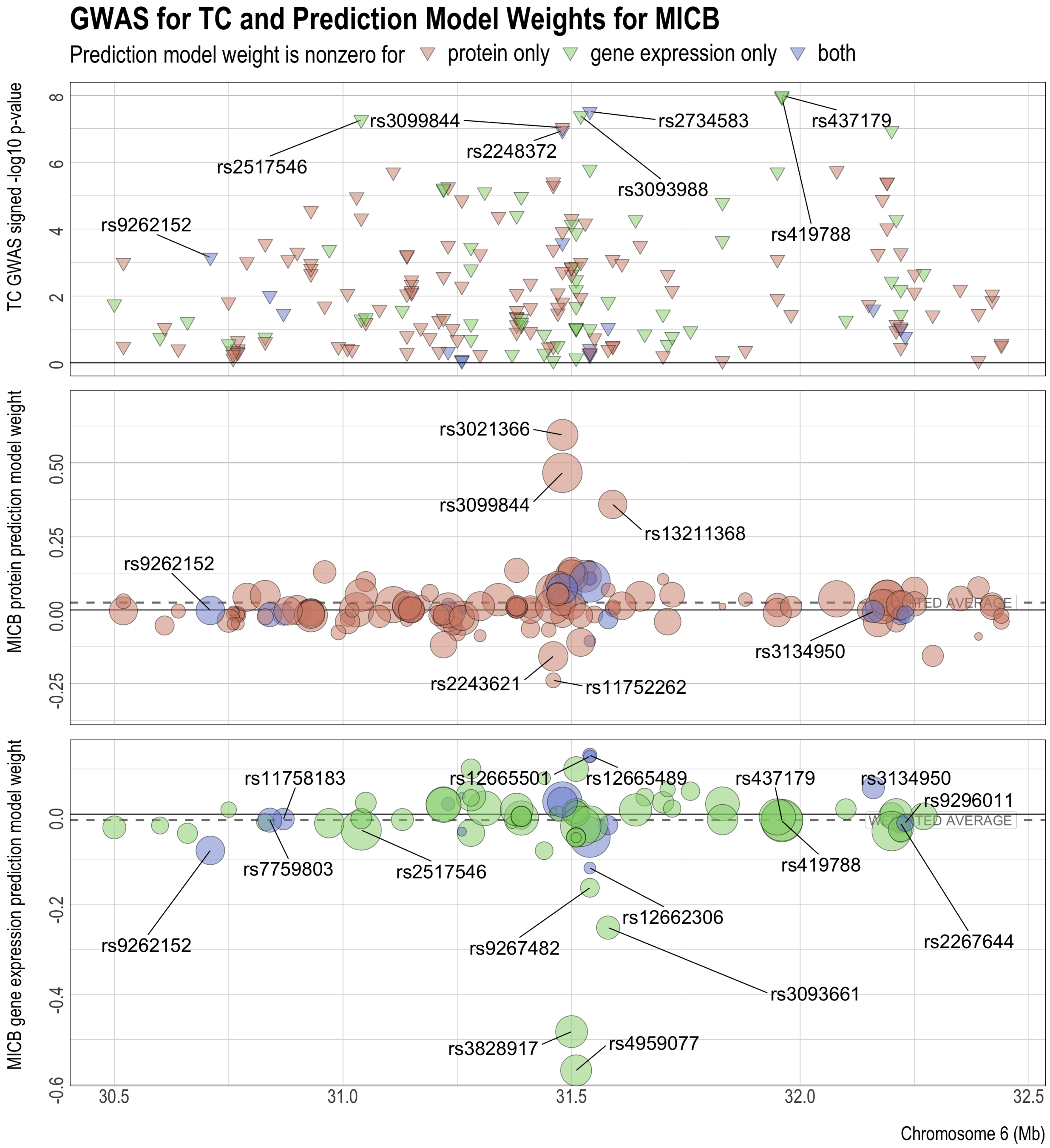

Figure S16: Comparison of MICB's protein and gene expression predictive model weights with the TC GWAS z-scores of the SNPs. The reference and alternative alleles for GWAS and the predictive models have been aligned and reordered so that all the SNPs have positive GWAS effects. The z-scores are used to compute the weighted average of the model weights (dashed lines), which have the same signs as and are proportional to the predicted effects of protein and gene expression on the GWAS outcome.

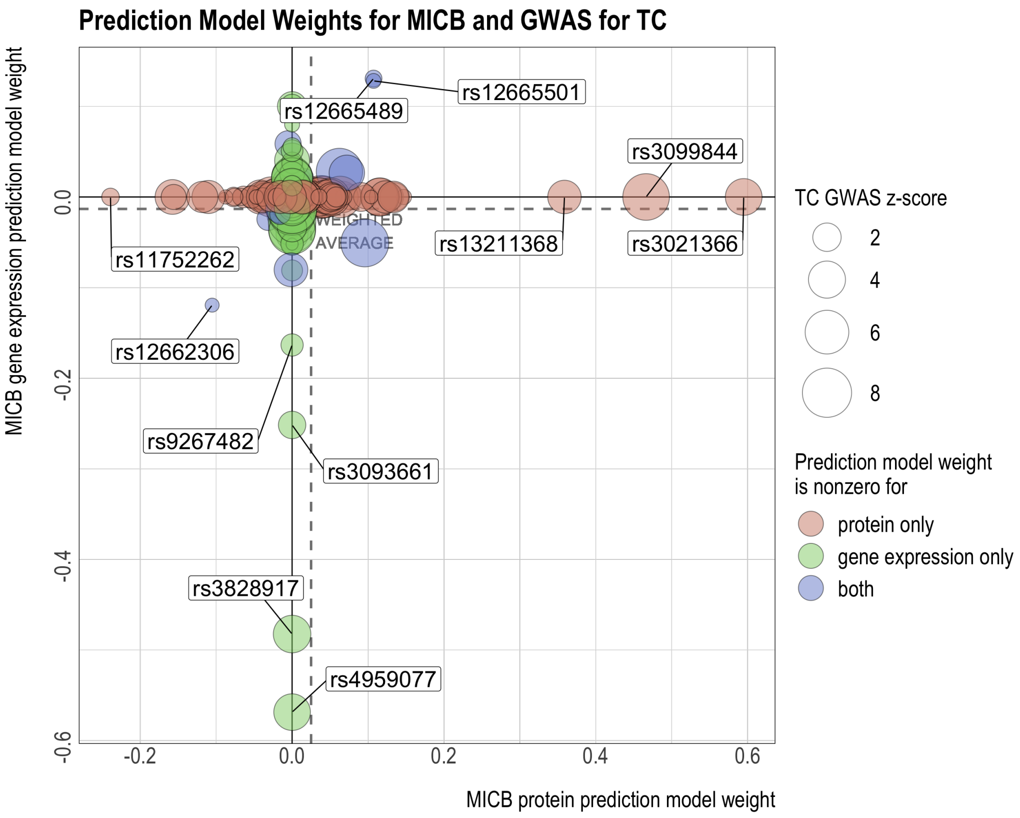

Figure S17: Comparison of MESA PBMC PWAS, MESA PBMC TWAS, and GTEx tissue-specific TWAS results for TC. Panel (a): signed log p-value and significance of association. Missing values are shown in white. Significance of association is deteremined by the false discovery rate (FDR) threshold of 0.05. Panel (b): correlation between signed log p-values of MESA PBMC PWAS and signed log p-values of each GTEx tissue-specific TWAS (i.e. the correlation between the bottom row and every other row of the grid in Panel (a)).

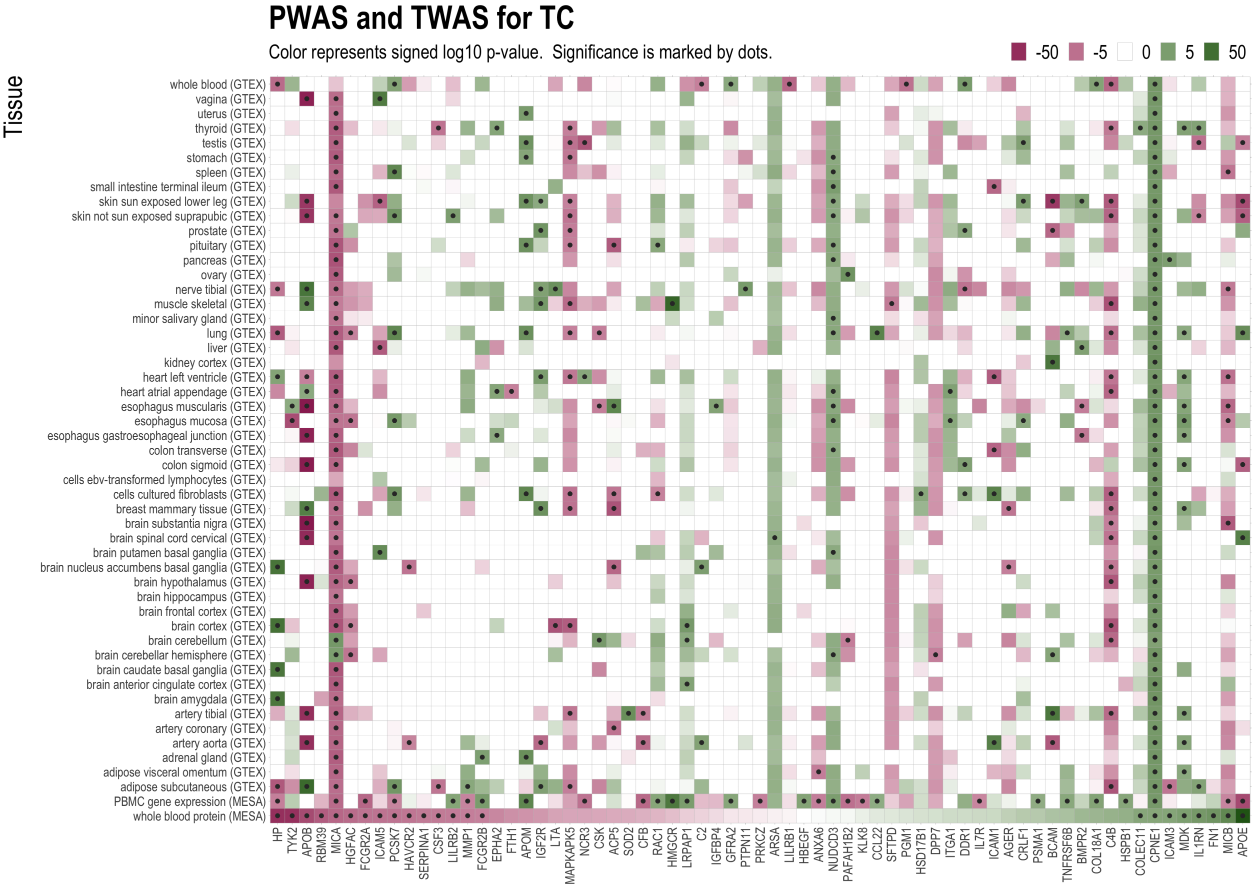


(a)

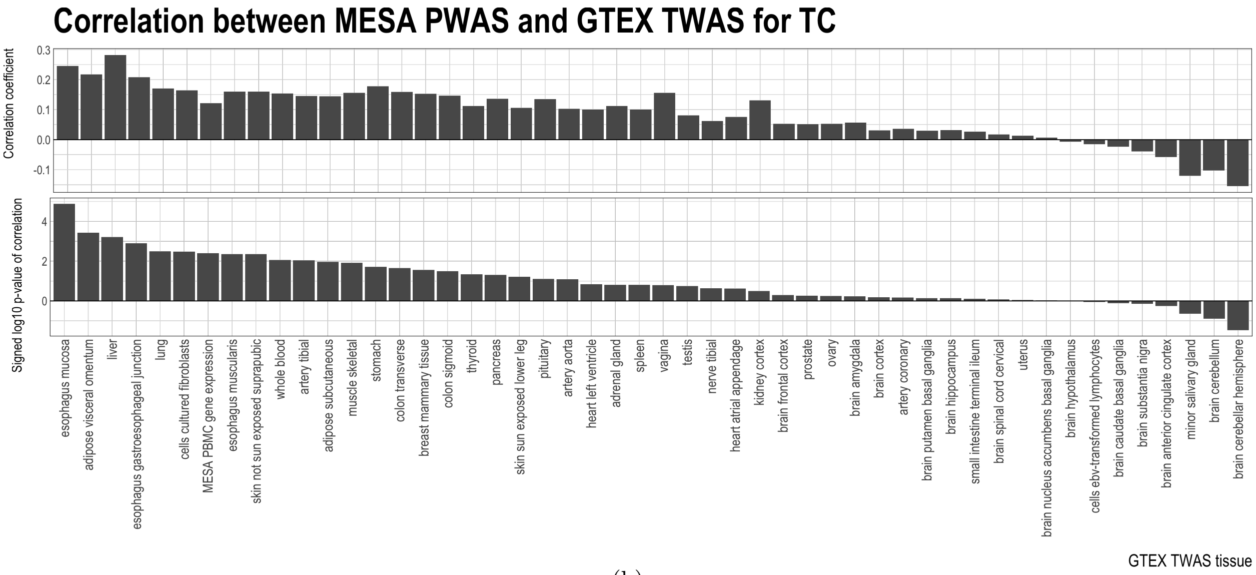


(b)

### S4 Additional results for triglycerides (TG)

Figure S18: GWAS for TG and prediction models for APOE's protein and gene expression levels. The reference and alternative alleles for GWAS and the predictive models have been aligned and reordered so that all the SNPs have positive GWAS effects. In the center and bottom panels, the size of the circles indicates the SNP's GWAS z-score. The z-scores are used to compute the weighted average of the model weights (dashed line), which has the same sign as and is proportional to the predicted effect of protein or gene expression on the GWAS outcome.

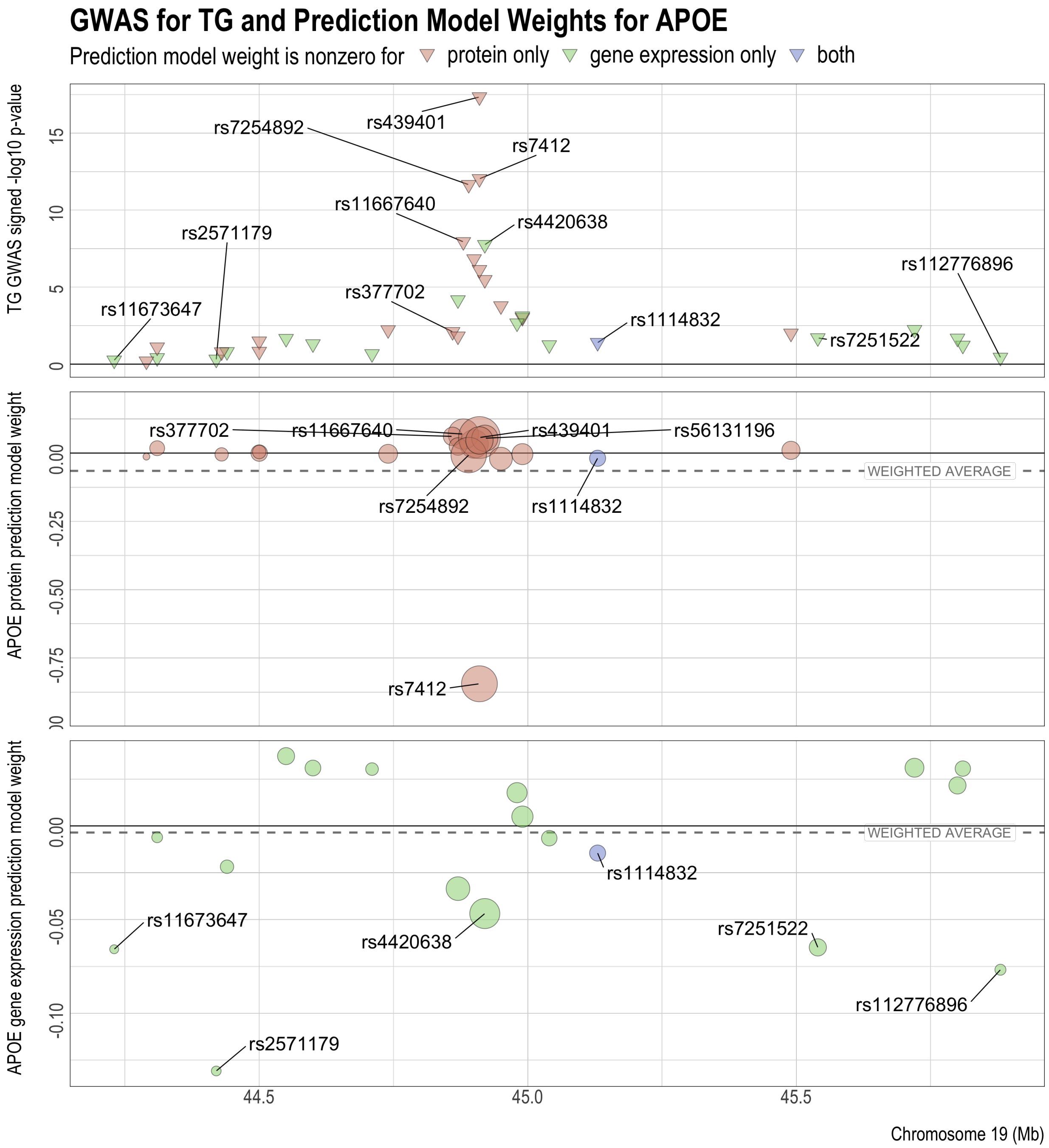

Figure S19: Comparison of APOE's protein and gene expression predictive model weights with the TG GWAS z-scores of the SNPs. The reference and alternative alleles for GWAS and the predictive models have been aligned and reordered so that all the SNPs have positive GWAS effects. The z-scores are used to compute the weighted average of the model weights (dashed lines), which have the same signs as and are proportional to the predicted effects of protein and gene expression on the GWAS outcome.

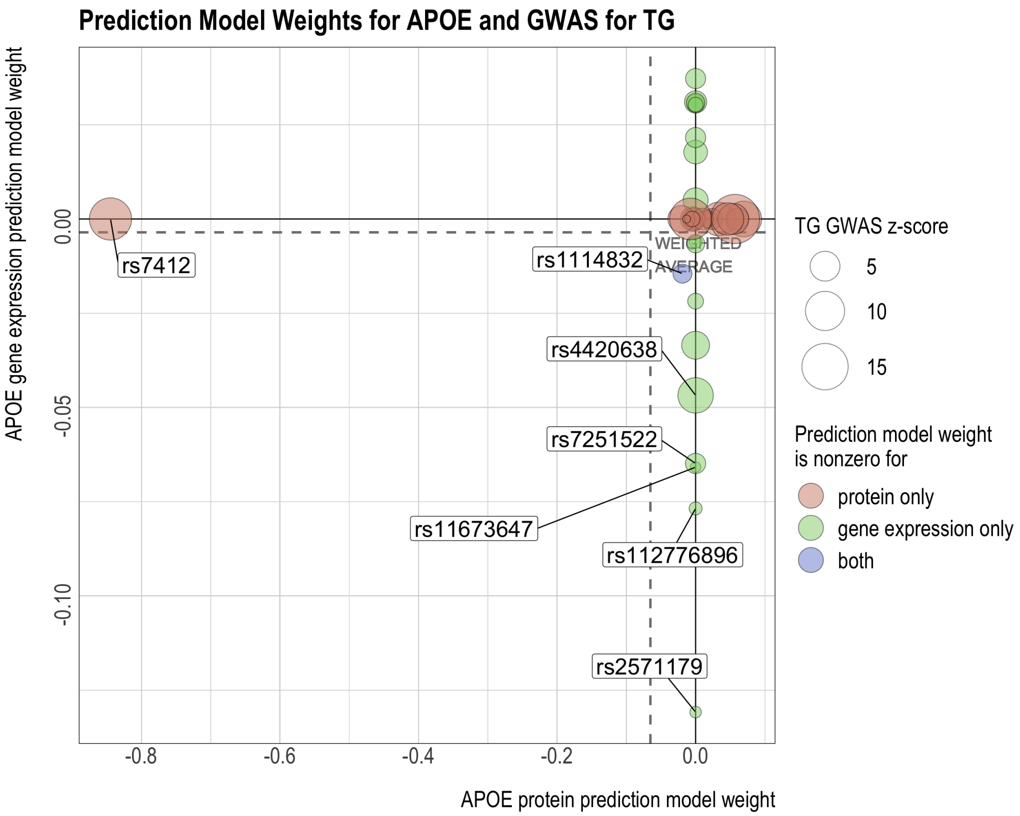

Figure S20: GWAS for TG and prediction models for FCGR2B's protein and gene expression levels. The reference and alternative alleles for GWAS and the predictive models have been aligned and reordered so that all the SNPs have positive GWAS effects. In the center and bottom panels, the size of the circles indicates the SNP's GWAS z-score. The z-scores are used to compute the weighted average of the model weights (dashed line), which has the same sign as and is proportional to the predicted effect of protein or gene expression on the GWAS outcome.

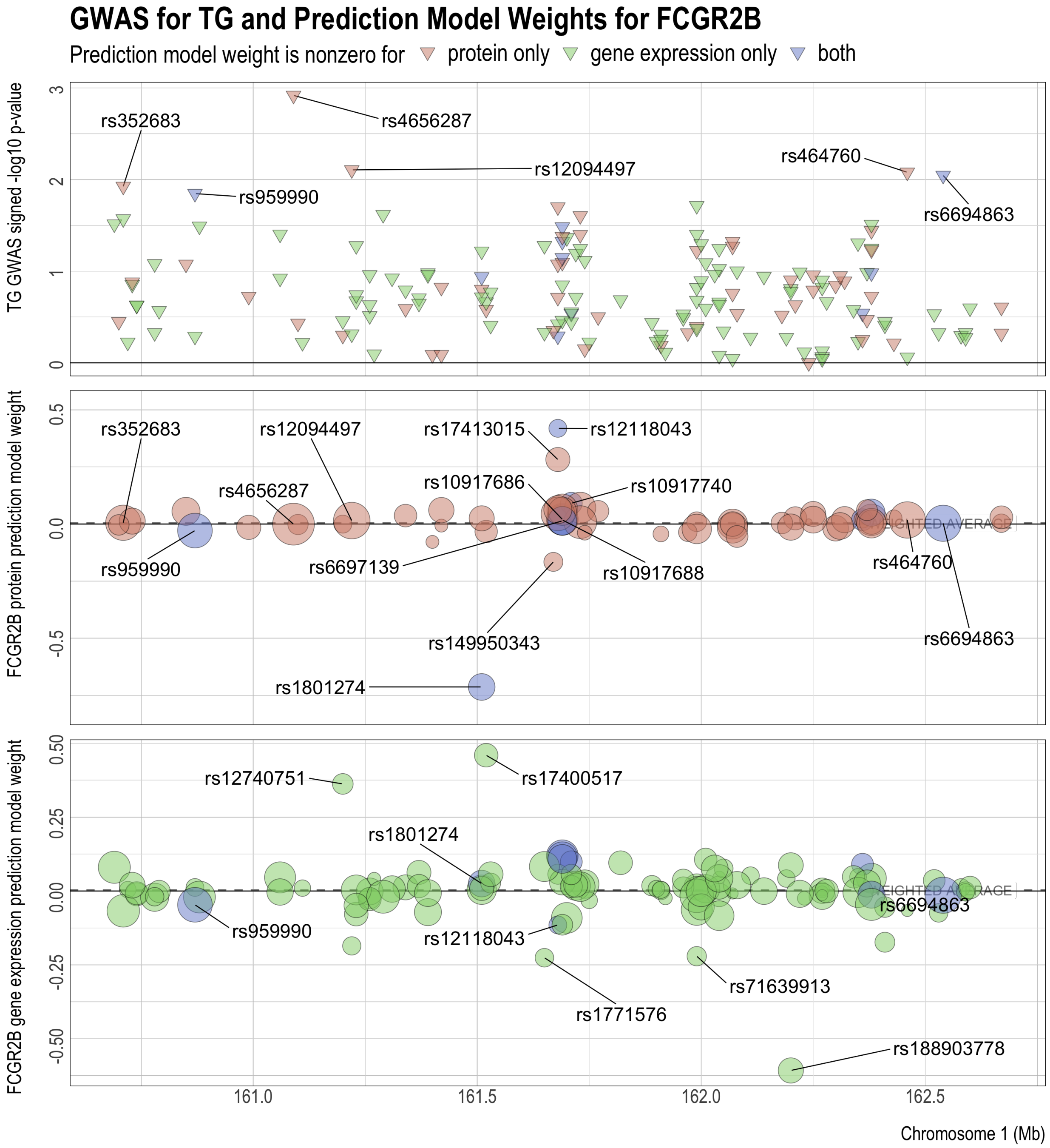

Figure S21: Comparison of FCGR2B's protein and gene expression predictive model weights with the TG GWAS z-scores of the SNPs. The reference and alternative alleles for GWAS and the predictive models have been aligned and reordered so that all the SNPs have positive GWAS effects. The z-scores are used to compute the weighted average of the model weights (dashed lines), which have the same signs as and are proportional to the predicted effects of protein and gene expression on the GWAS outcome.

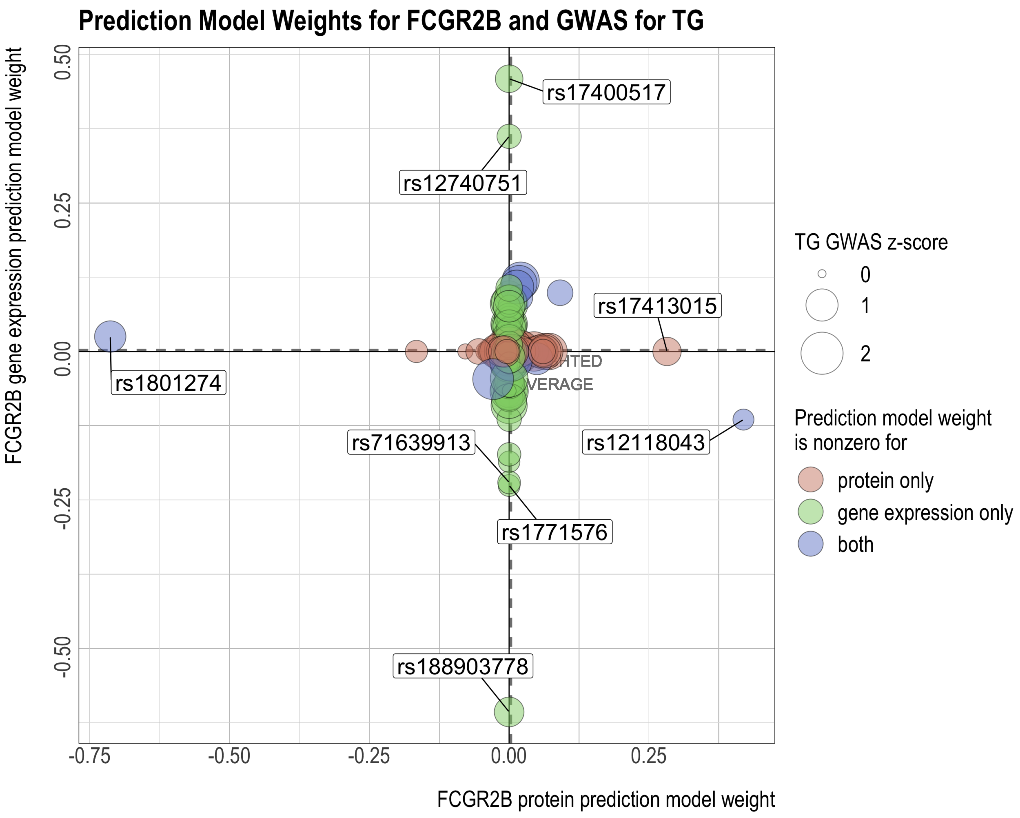

Figure S22: GWAS for TG and prediction models for LILRB2's protein and gene expression levels. The reference and alternative alleles for GWAS and the predictive models have been aligned and reordered so that all the SNPs have positive GWAS effects. In the center and bottom panels, the size of the circles indicates the SNP's GWAS z-score. The z-scores are used to compute the weighted average of the model weights (dashed line), which has the same sign as and is proportional to the predicted effect of protein or gene expression on the GWAS outcome.

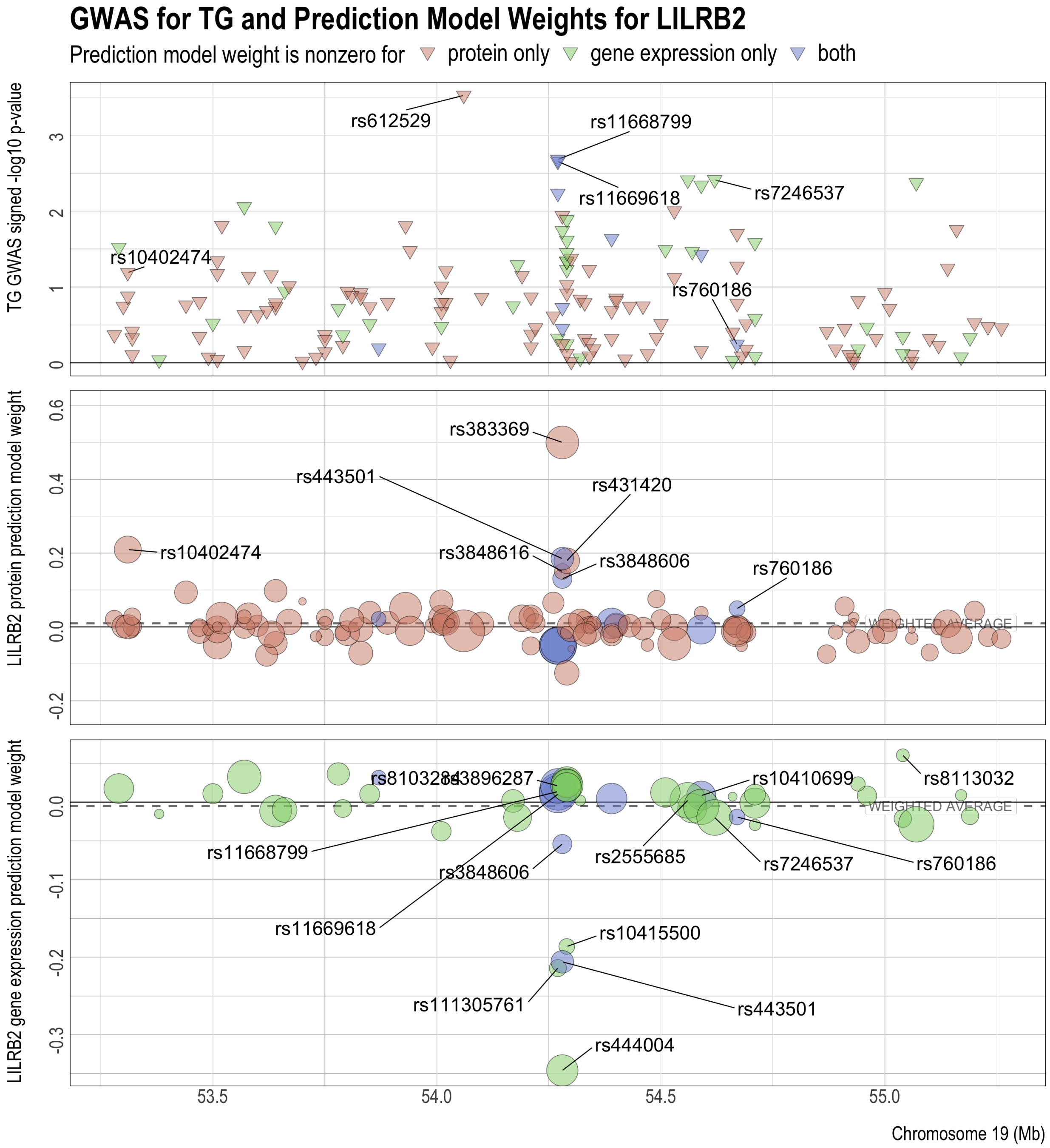

Figure S23: Comparison of LILRB2's protein and gene expression predictive model weights with the TG GWAS z-scores of the SNPs. The reference and alternative alleles for GWAS and the predictive models have been aligned and reordered so that all the SNPs have positive GWAS effects. The z-scores are used to compute the weighted average of the model weights (dashed lines), which have the same signs as and are proportional to the predicted effects of protein and gene expression on the GWAS outcome.

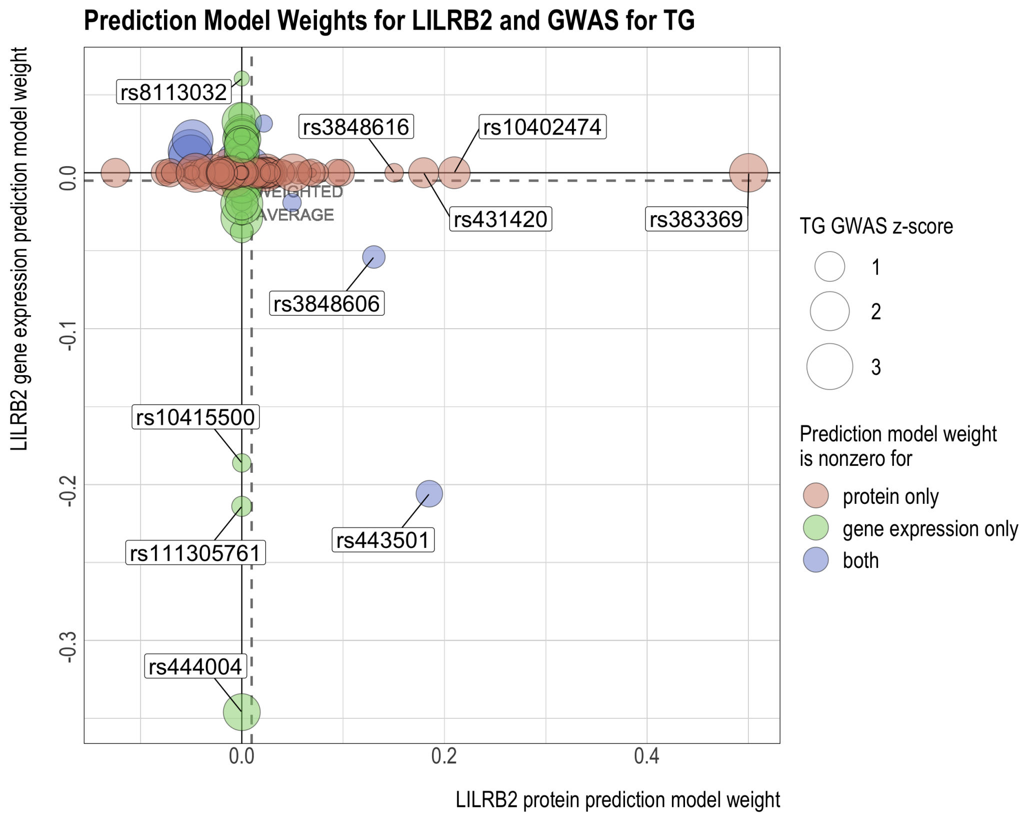

Figure S24: GWAS for TG and prediction models for MICB's protein and gene expression levels. The reference and alternative alleles for GWAS and the predictive models have been aligned and reordered so that all the SNPs have positive GWAS effects. In the center and bottom panels, the size of the circles indicates the SNP's GWAS z-score. The z-scores are used to compute the weighted average of the model weights (dashed line), which has the same sign as and is proportional to the predicted effect of protein or gene expression on the GWAS outcome.

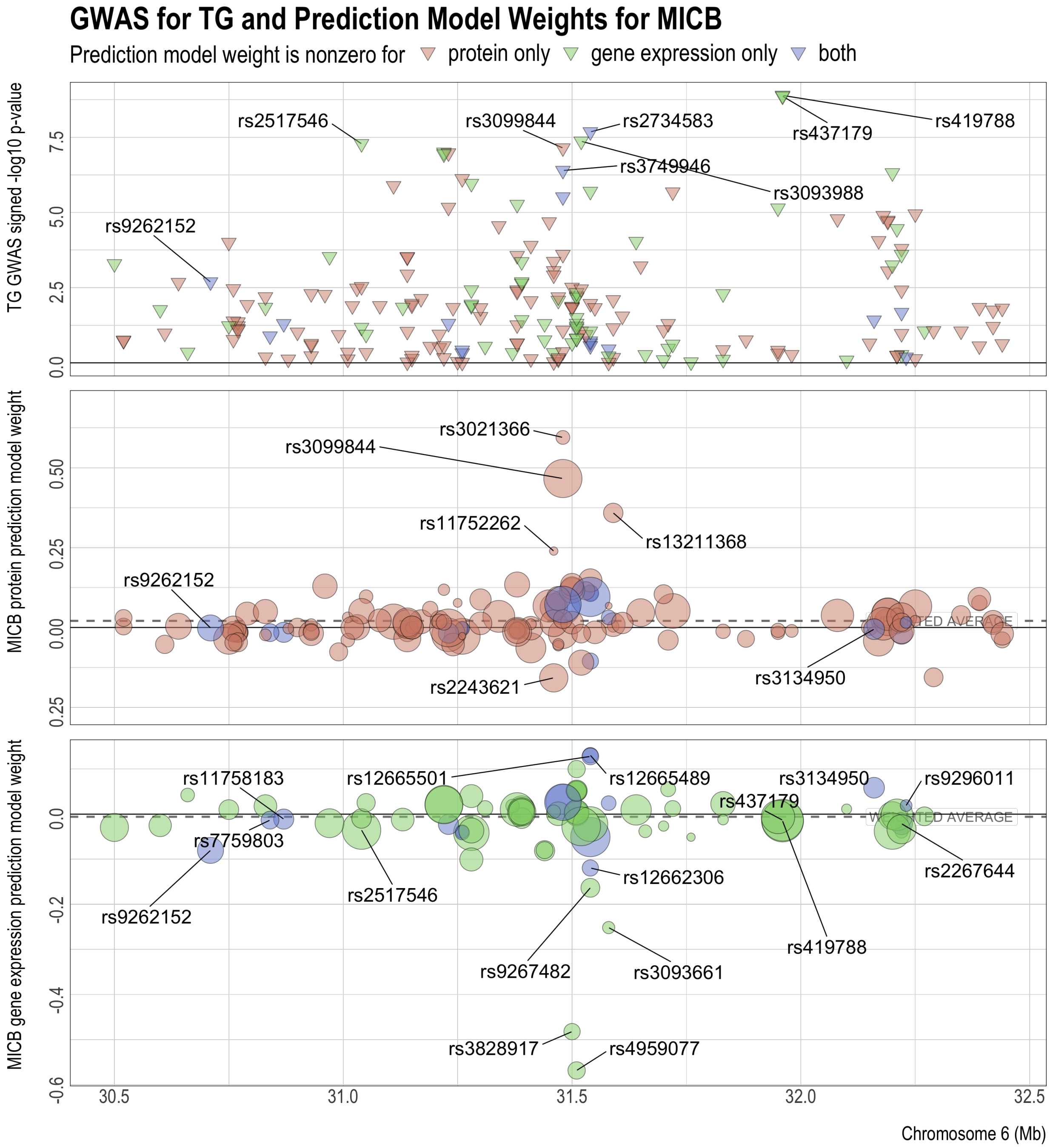

Figure S25: Comparison of MICB's protein and gene expression predictive model weights with the TG GWAS z-scores of the SNPs. The reference and alternative alleles for GWAS and the predictive models have been aligned and reordered so that all the SNPs have positive GWAS effects. The z-scores are used to compute the weighted average of the model weights (dashed lines), which have the same signs as and are proportional to the predicted effects of protein and gene expression on the GWAS outcome.

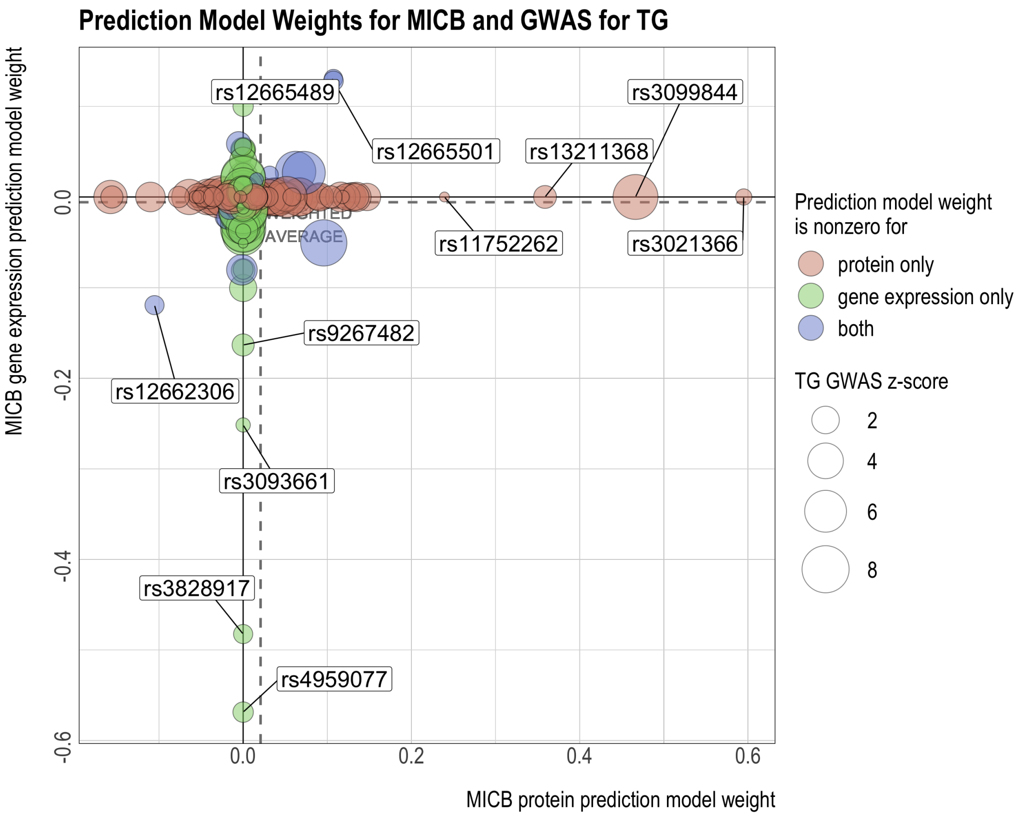

Figure S26: Comparison of MESA PBMC PWAS, MESA PBMC TWAS, and GTEx tissue-specific TWAS results for TG. Panel (a): signed log p-value and significance of association. Missing values are shown in white. Significance of association is deteremined by the false discovery rate (FDR) threshold of 0.05. Panel (b): correlation between signed log p-values of MESA PBMC PWAS and signed log p-values of each GTEx tissue-specific TWAS (i.e. the correlation between the bottom row and every other row of the grid in Panel (a)).

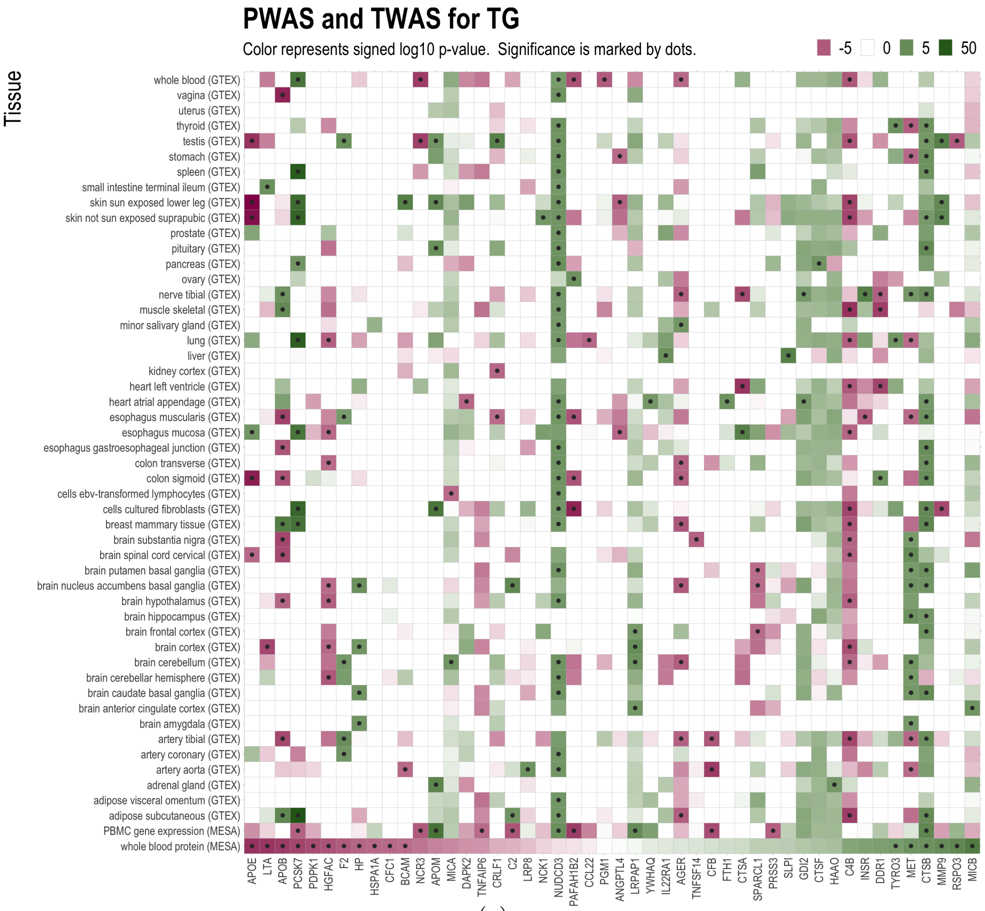


(a)

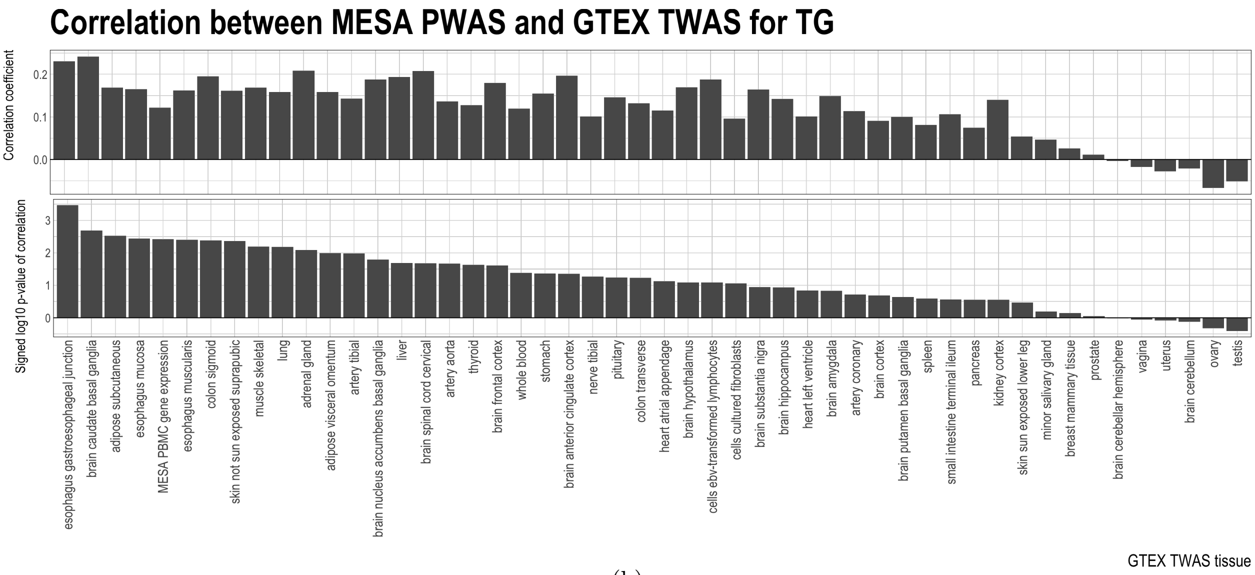


(b)

### S5 Additional results for high-density lipoprotein (HDL)

Figure S27: GWAS for HDL and prediction models for APOE's protein and gene expression levels. The reference and alternative alleles for GWAS and the predictive models have been aligned and reordered so that all the SNPs have positive GWAS effects. In the center and bottom panels, the size of the circles indicates the SNP's GWAS z-score. The z-scores are used to compute the weighted average of the model weights (dashed line), which has the same sign as and is proportional to the predicted effect of protein or gene expression on the GWAS outcome.

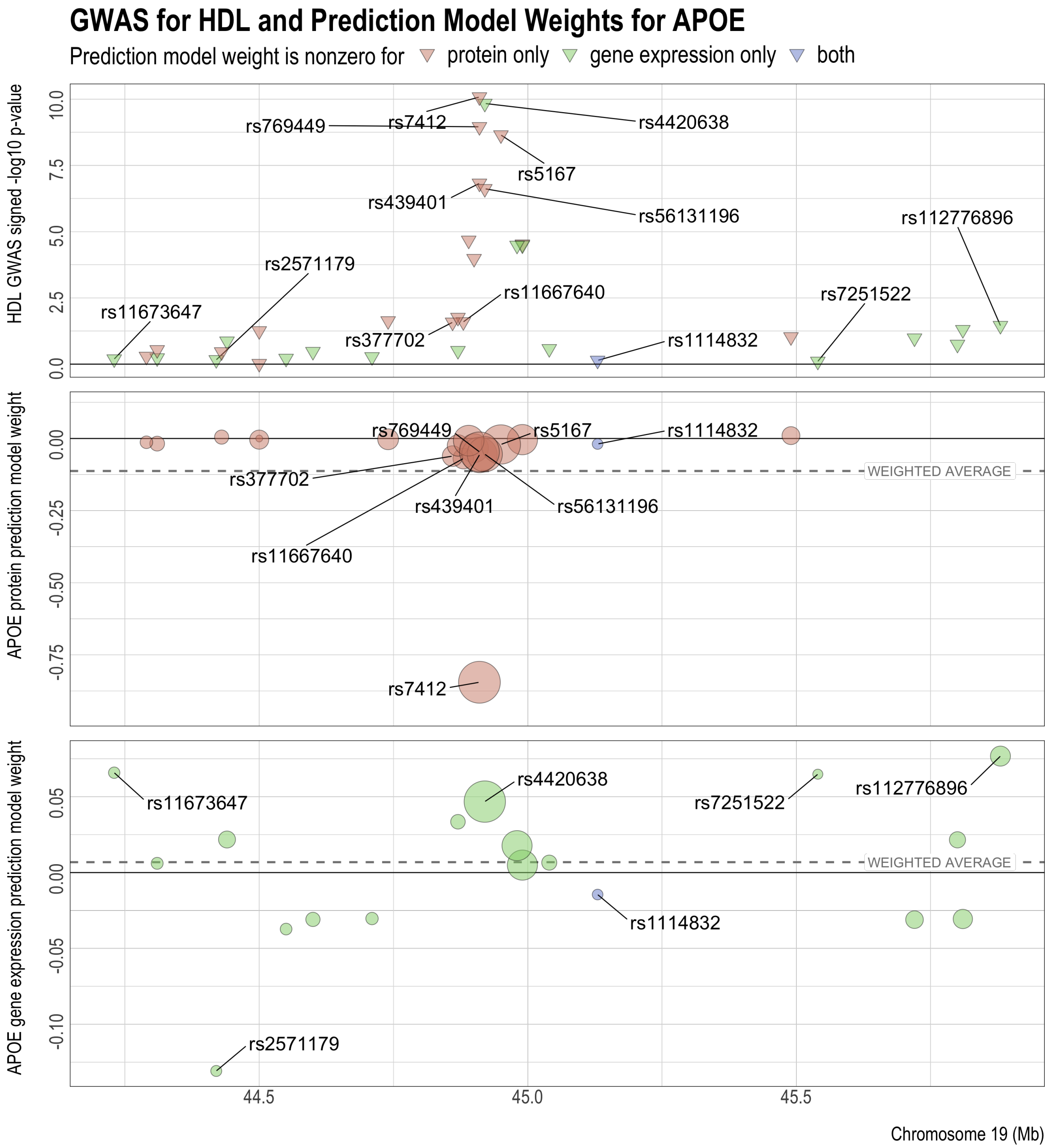

Figure S28: Comparison of APOE's protein and gene expression predictive model weights with the HDL GWAS z-scores of the SNPs. The reference and alternative alleles for GWAS and the predictive models have been aligned and reordered so that all the SNPs have positive GWAS effects. The z-scores are used to compute the weighted average of the model weights (dashed lines), which have the same signs as and are proportional to the predicted effects of protein and gene expression on the GWAS outcome.

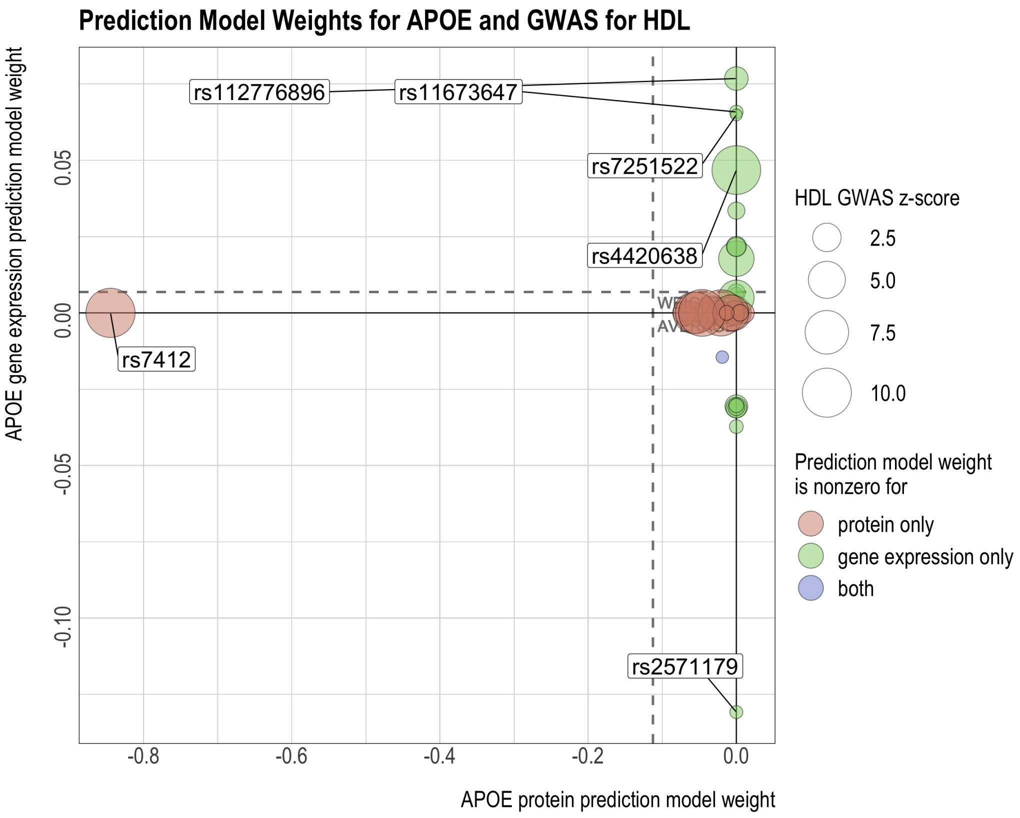

Figure S29: GWAS for HDL and prediction models for FCGR2B's protein and gene expression levels. The reference and alternative alleles for GWAS and the predictive models have been aligned and reordered so that all the SNPs have positive GWAS effects. In the center and bottom panels, the size of the circles indicates the SNP's GWAS z-score. The z-scores are used to compute the weighted average of the model weights (dashed line), which has the same sign as and is proportional to the predicted effect of protein or gene expression on the GWAS outcome.

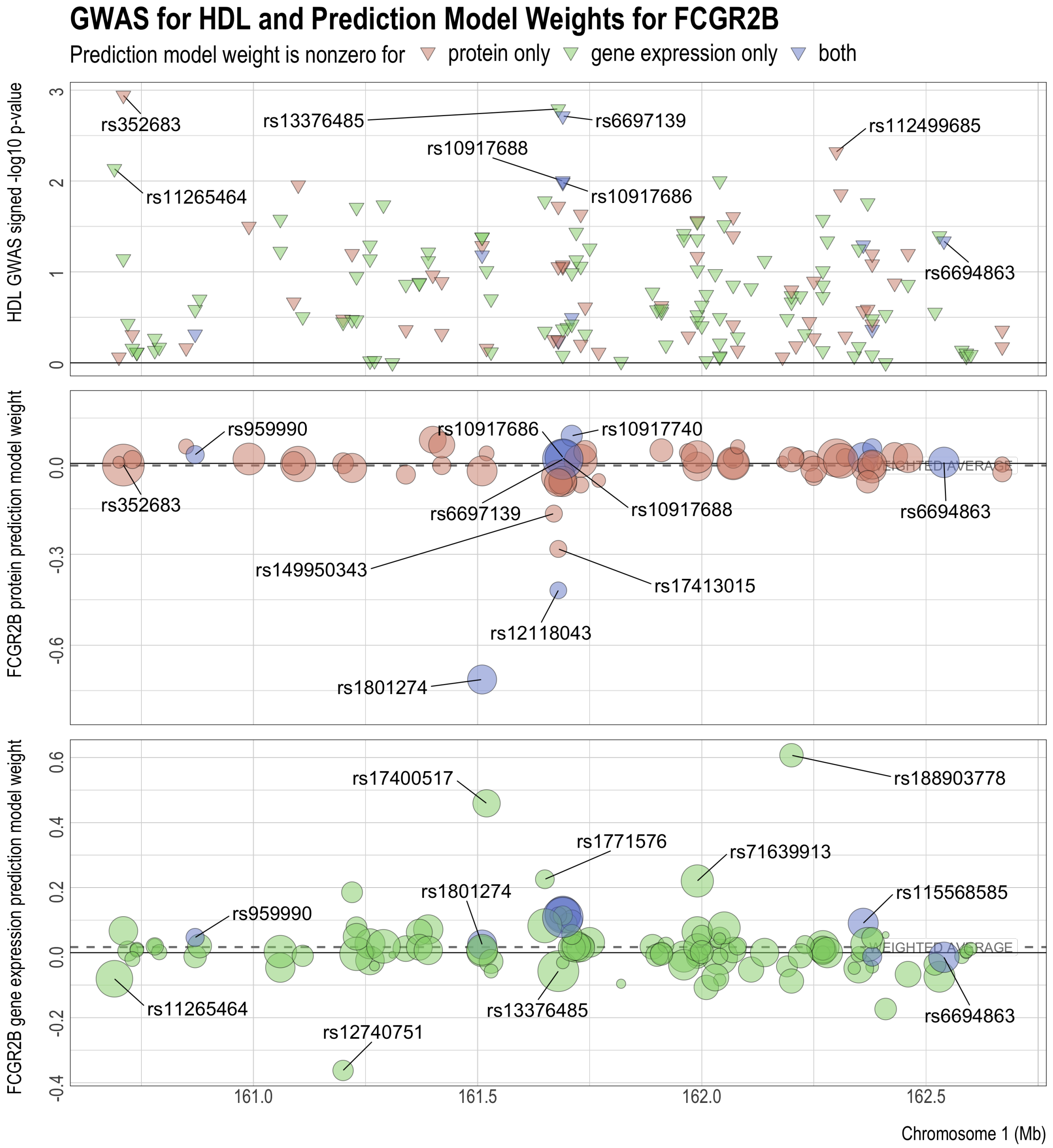

Figure S30: Comparison of FCGR2B's protein and gene expression predictive model weights with the HDL GWAS z-scores of the SNPs. The reference and alternative alleles for GWAS and the predictive models have been aligned and reordered so that all the SNPs have positive GWAS effects. The z-scores are used to compute the weighted average of the model weights (dashed lines), which have the same signs as and are proportional to the predicted effects of protein and gene expression on the GWAS outcome.

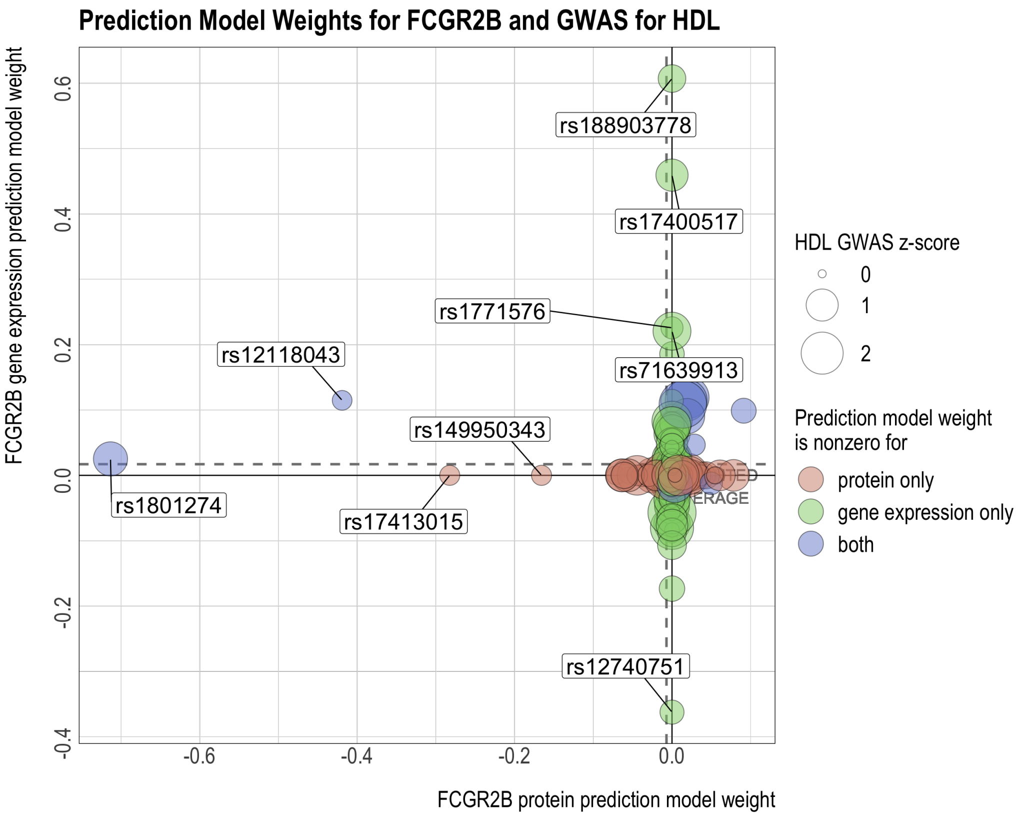

Figure S31: GWAS for HDL and prediction models for LILRB2's protein and gene expression levels. The reference and alternative alleles for GWAS and the predictive models have been aligned and reordered so that all the SNPs have positive GWAS effects. In the center and bottom panels, the size of the circles indicates the SNP's GWAS z-score. The z-scores are used to compute the weighted average of the model weights (dashed line), which has the same sign as and is proportional to the predicted effect of protein or gene expression on the GWAS outcome.

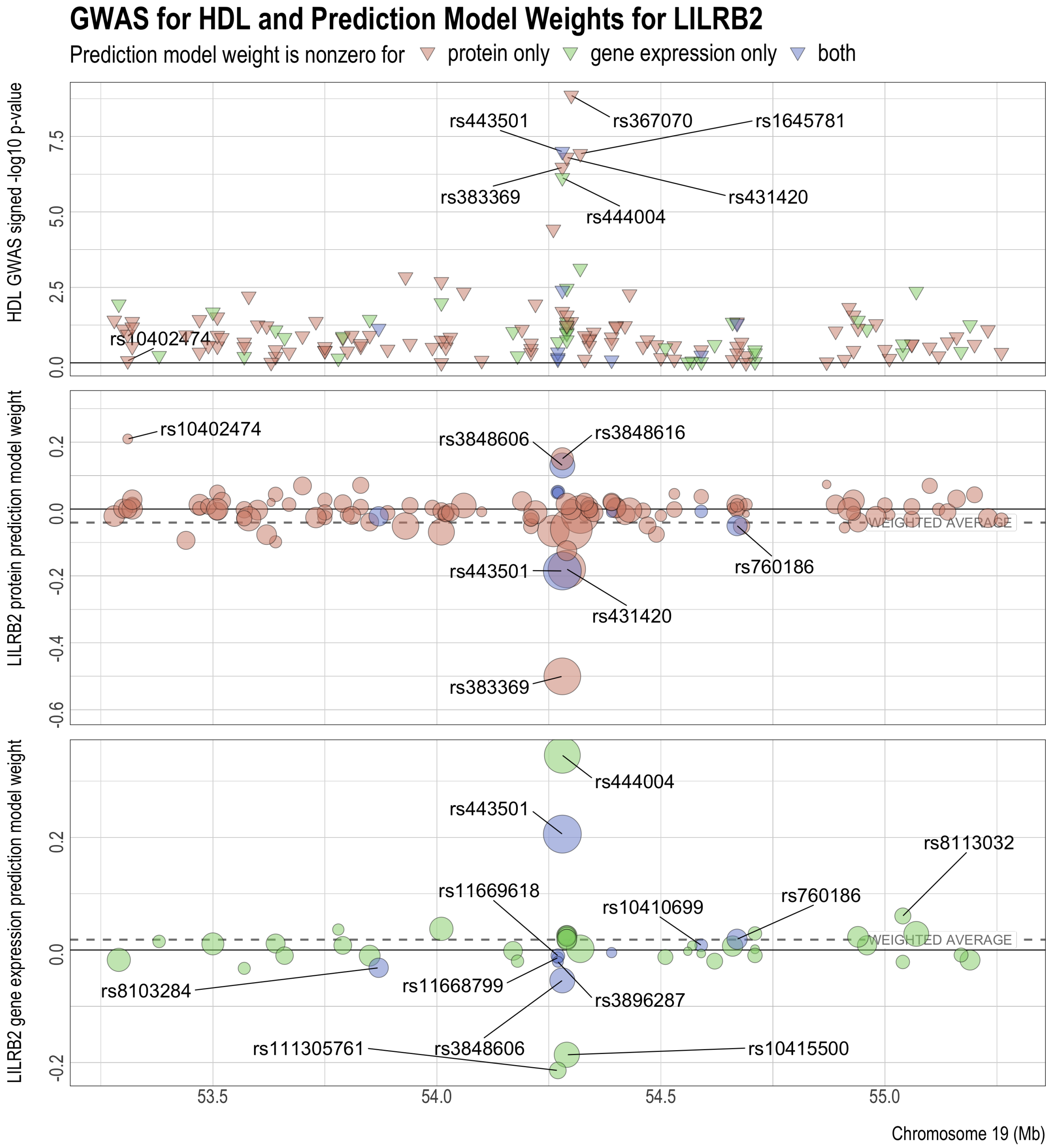

Figure S32: Comparison of LILRB2's protein and gene expression predictive model weights with the HDL GWAS z-scores of the SNPs. The reference and alternative alleles for GWAS and the predictive models have been aligned and reordered so that all the SNPs have positive GWAS effects. The z-scores are used to compute the weighted average of the model weights (dashed lines), which have the same signs as and are proportional to the predicted effects of protein and gene expression on the GWAS outcome.

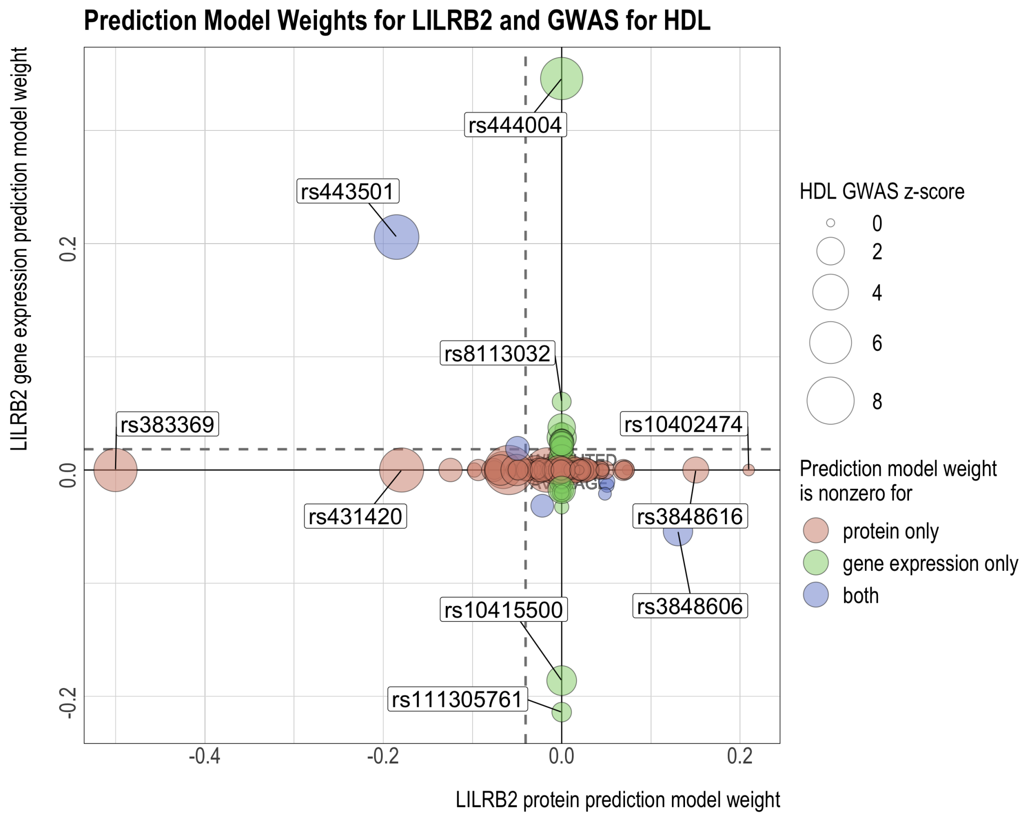

Figure S33: GWAS for HDL and prediction models for MICB's protein and gene expression levels. The reference and alternative alleles for GWAS and the predictive models have been aligned and reordered so that all the SNPs have positive GWAS effects. In the center and bottom panels, the size of the circles indicates the SNP's GWAS z-score. The z-scores are used to compute the weighted average of the model weights (dashed line), which has the same sign as and is proportional to the predicted effect of protein or gene expression on the GWAS outcome.

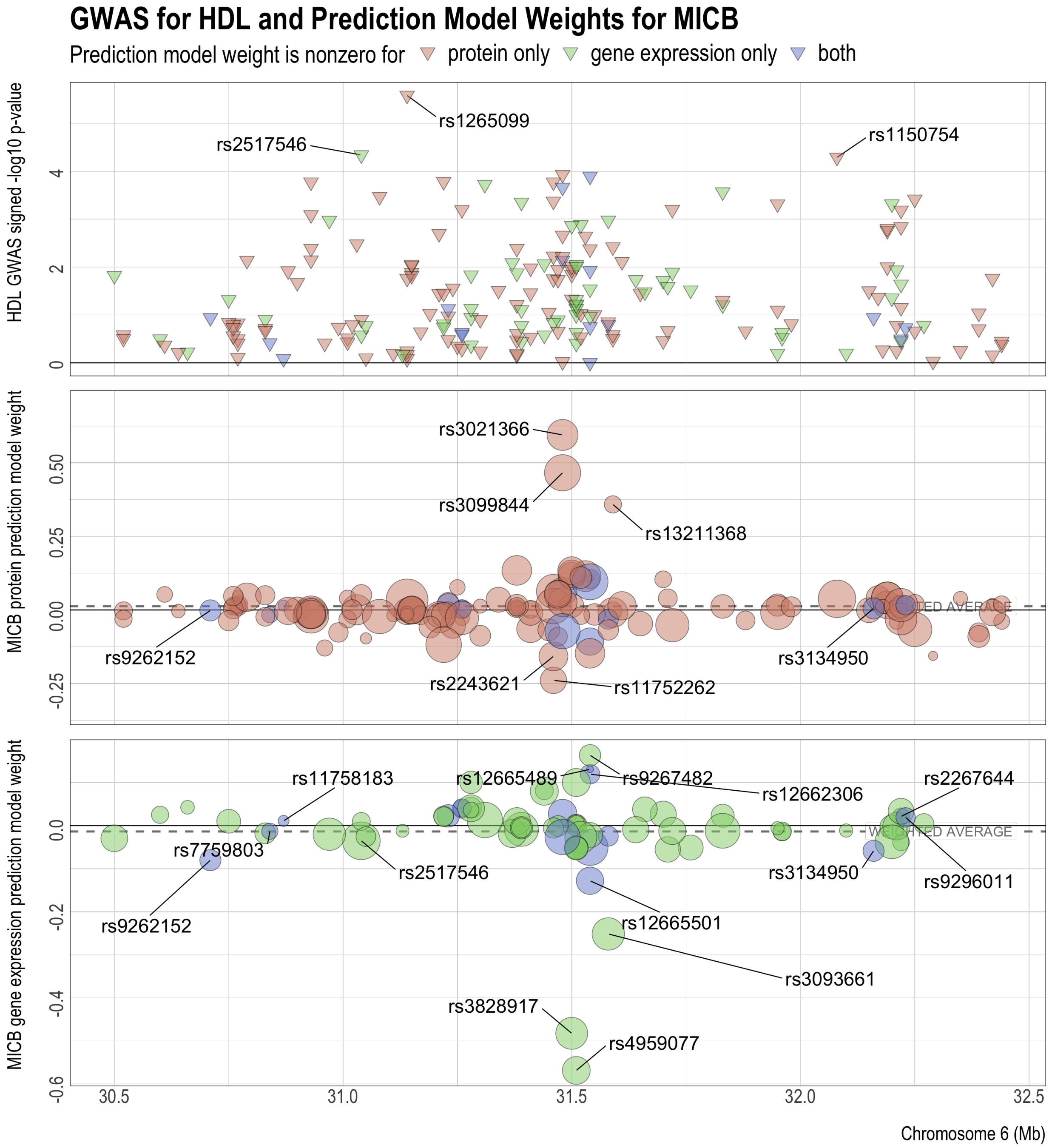

Figure S34: Comparison of MICB's protein and gene expression predictive model weights with the HDL GWAS z-scores of the SNPs. The reference and alternative alleles for GWAS and the predictive models have been aligned and reordered so that all the SNPs have positive GWAS effects. The z-scores are used to compute the weighted average of the model weights (dashed lines), which have the same signs as and are proportional to the predicted effects of protein and gene expression on the GWAS outcome.

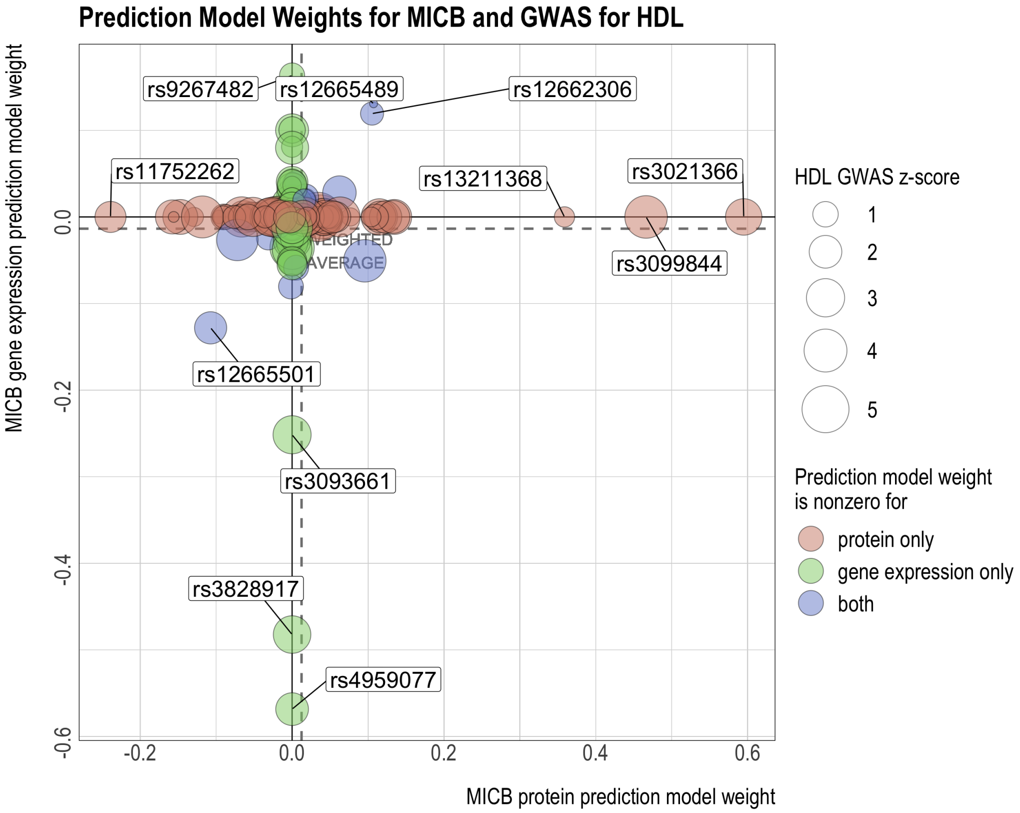

Figure S35: Comparison of MESA PBMC PWAS, MESA PBMC TWAS, and GTEx tissue-specific TWAS results for HDL. Panel (a): signed log p-value and significance of association. Missing values are shown in white. Significance of association is deteremined by the false discovery rate (FDR) threshold of 0.05. Panel (b): correlation between signed log p-values of MESA PBMC PWAS and signed log p-values of each GTEx tissue-specific TWAS (i.e. the correlation between the bottom row and every other row of the grid in Panel (a)).

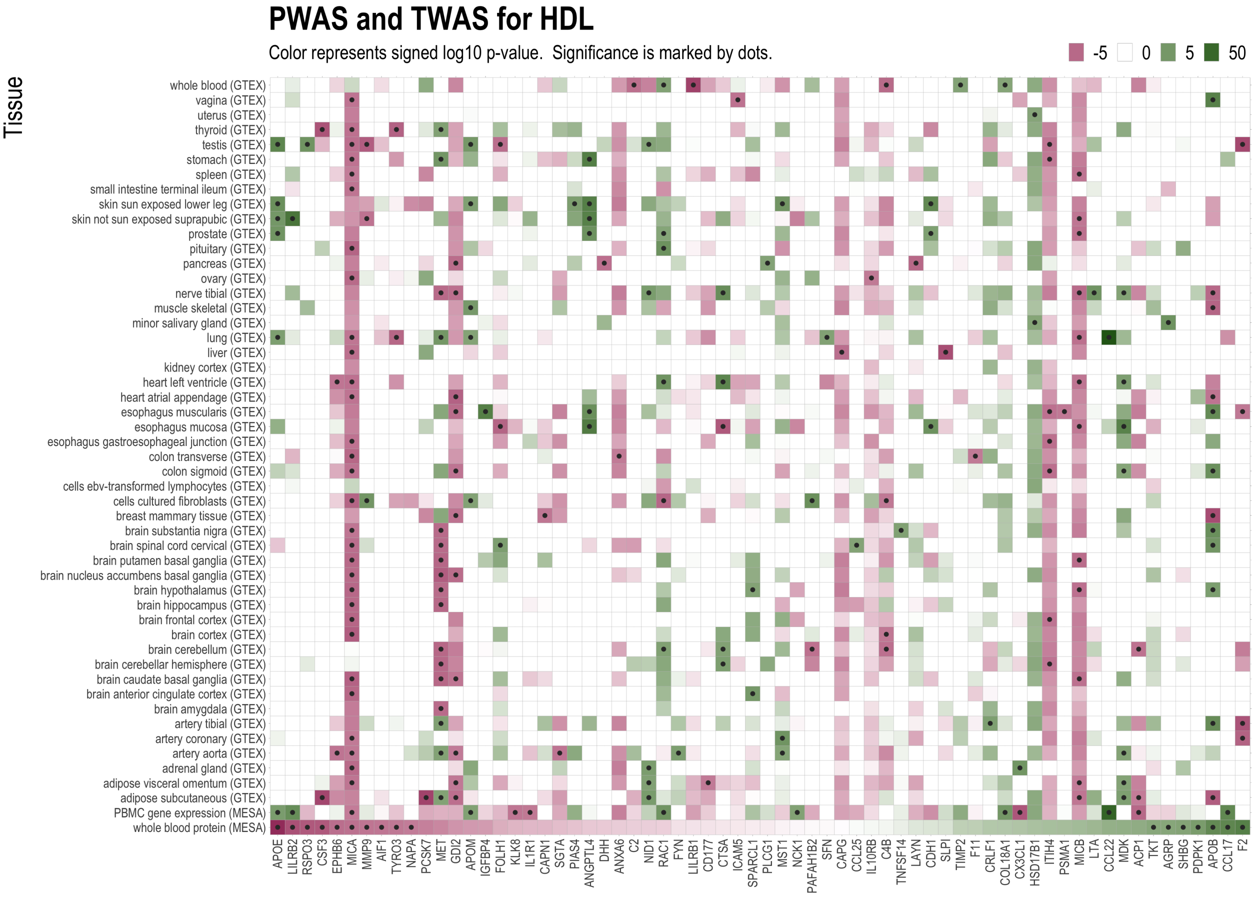


(a)

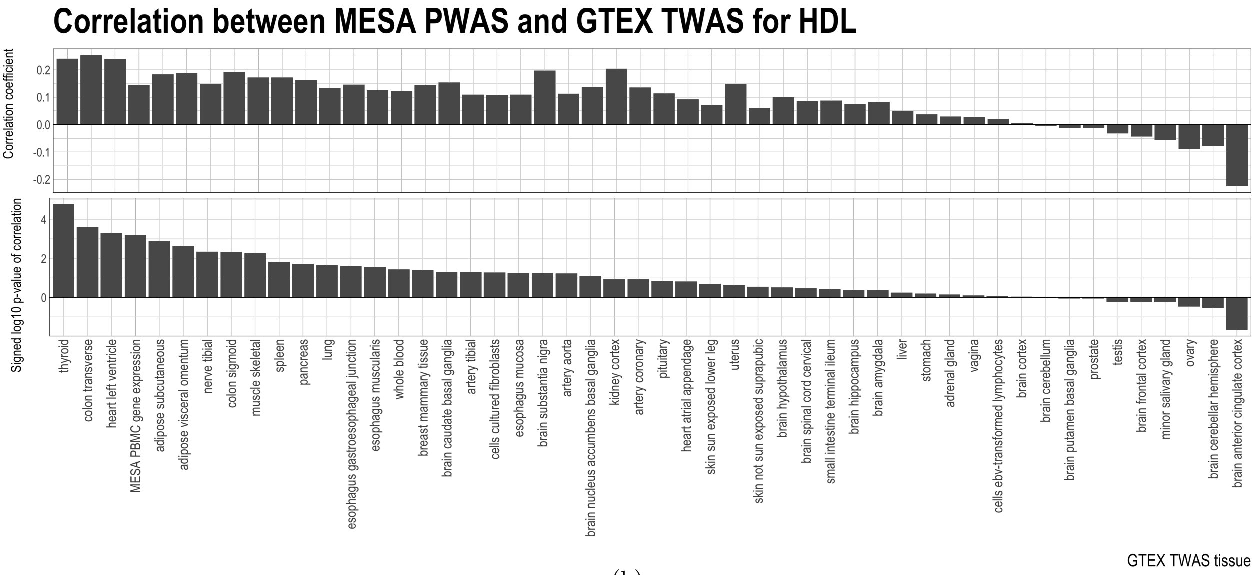


(b)
